# Haplotype analyses reveal novel insights into tomato history and domestication including long-distance migrations and latitudinal adaptations

**DOI:** 10.1101/2021.06.18.448912

**Authors:** Jose Blanca, David Sanchez-Matarredona, Peio Ziarsolo, J Montero-Pau, Esther van der Knaap, Maria José Díez, Joaquín Cañizares

**Author notes:** Author for correspondence: Jose Blanca.

## Abstract

A novel haplotype-based approach that uses Procrustes analysis and automatic classification was used to provide further insights into tomato history and domestication. Agrarian societies domesticated species of interest by introducing complex genetic modifications. For tomatoes, two species, one of which had two botanical varieties, are thought to be involved in its domestication: the fully wild *Solanum pimpinellifolium* (SP), the wild and semi-domesticated *S. lycopersicum* var. *cerasiforme* (SLC) and the cultivated *S. l.* var. *lycopersicum* (SLL). The Procrustes approach showed that SP evolved into SLC during a gradual migration from the Peruvian deserts to the Mexican rainforests and that Peruvian and Ecuadorian SLC populations were the result of more recent hybridizations. Our model was supported by independent evidence, including ecological data from the accession collection site and morphological data. Furthermore, we showed that photosynthesis-, and flowering time-related genes were selected during the latitudinal migrations.

## Introduction

Cultivated plants result from domestication processes that alter the morphology, physiology, and genetics of wild species to benefit human needs and preferences. These processes usually involve a domestication syndrome, which involves modifying a set of traits (Hammer, 1984) favored by humans, and/or that provide growth advantages under cultivation or adaptations to thrive in disturbed habitats (Meyer & Purugganan, 2013). In horticultural crops, these traits usually include larger and more nutritious fruits, robust stems, and reduced seed dormancy (Yang, Li, Tieman, & Zhu, 2019). Selection, bottlenecks, and outcrossing with wild and feral populations are common during domestication. Moreover, domestication histories are intertwined with the history of the agrarian cultures that performed them, and complex migrations and interchanges between different geographic regions have also occurred. Thus, the extant population genetic structure and the patterns of morphological diversity are often complex. In addition to historical interest, the study of these processes has practical implications because they generate knowledge regarding genes and pathways of agronomic interest (Zsögön et al., 2018).

The fully wild *Solanum pimpinellifolium* L. (SP) and *S. lycopersicum* L. (SL) are two sister Solanaceae species (genus *Solanum* L., section *Lycopersicon* (Peralta, Spooner, & Knapp, 2008), which are capable of interbreeding. SL is split into two botanical varieties: *S. l.* var. *lycopersicum* L. (SLL) and *S. l.* var. *cerasiforme* (Dunal) Spooner, G.J. Anderson & R.K. Jansen (SLC) (Peralta et al., 2008). SP has been proposed as the species from which the cultivated tomato forms have been domesticated (Peralta et al., 2008). SP is divided into several populations associated with different climates and ecological niches: the dry Peruvian coast, the northern Peruvian and southern Ecuadorian Andean valleys, and the wet northern Ecuadorian coast (Gibson & Moyle, 2020).

Recent studies have shown that SP likely evolved into SLC before human colonization of the Americas (Razifard et al., 2020). Therefore, an ancestral wild SLC population might have been involved in tomato domestication. SLL is cultivated, whereas SLC comprises a complex mix of wild, semi-domesticated, and vintage Peruvian, Ecuadorian, and Mesoamerican varieties (José Blanca et al., 2015; C. M. Rick & Holle, 1990). Furthermore, as a feral and weedy species, SLC colonized subtropical regions worldwide after the arrival of the Europeans in America (C. M. Rick & Holle, 1990).

The most accepted model for tomato domestication is a two-step process (Jose Blanca et al., 2012; José Blanca et al., 2015; Gao et al., 2019; Lin et al., 2014; Razifard et al., 2020). According to this model, the desert-dwelling, most diverse, and wild Peruvian SP (SP Pe) population comprised the most ancient population. In a slow process unrelated to human activity, SP Pe adapted to the climatic conditions in the Peruvian and Ecuadorian Andean valleys (SP Montane) and the northern humid Ecuadorian regions (SP Ec). Ecuadorian SLC (SLC Ec), which comprises the most diverse SLC population and has the shortest genetic distance to any SP, would be the first SL type derived from SP (Jose Blanca et al., 2012). SLC Ec occupies the humid Amazonian regions in Ecuador closer to the Andes, also known as Ceja de la Montaña (C. M. Rick & Holle, 1990). Afterward, SLC would have moved south from Ecuador to northern Peru, where early farmers might have begun domestication. In Ecuador and northern Peru, SLC accessions with a domesticated fruit morphology were found, and several Peruvian and Ecuadorian cultivated SLCs were collected in markets in the early part of the last century. Finally, the Peruvian cultivated tomatoes would have migrated to Mesoamerica, and there, in a second phase of improvement, SLL emerged.

However, there are complexities in tomato domestication history that remain to be explained (Jose Blanca et al., 2012; Razifard et al., 2020). For instance, SLC Ec is the most genetically diverse SLC, but it is proposed to be derived from a less diverse population, SP Ec. The conclusions of prior evolutionary genetic analyses were supported, at least partially, using traditional population indices, such as genetic diversity or linkage disequilibrium (LD). Despite being informative, these values can be misleading because they may be accounted by different hypotheses. For instance, high genetic diversity is typically found in ancient and well-established populations and recent admixtures. When several measures are combined, certain hypotheses can be effectively refuted. However, when the evolutionary history is complex, conclusions based on these traditional indices remain tentative. Complex statistical models may be used as alternatives to these nonparametric approaches. However, these complex parametric methods also have limitations because they tend to make unrealistic assumptions, such as the lack of Hardy-Weinberg violations, and/or depend on parameters that are difficult to establish, such as the number of ancestral migrations (Pickrell & Pritchard, 2012; Raj, Stephens, & Pritchard, 2014; Razifard et al., 2020).

Recently, Razifard et al. (2020) interpreted the high genetic diversity of SLC Ec by assuming that it was an ancient and wild population closely related to SP. However, their data are inconsistent with this hypothesis: SLC Ec appears to be an admixture according to their fastStructure results, and, although we would expect LD to be lower in the older population, SLC Ec had a higher LD than the Mexican SLC.

Linguistic and historical evidence might complement the genetic data in the study of domestication history. However, there is scant linguistic and historical evidence for tomato, and what is available is ambiguous and subject to different interpretations (Iris Peralta & David Spooner, 2011). Moreover, few tomato archeological remains have been uncovered. This might, at least in part, be caused by the perishability of the tomatoes (Kiple, Ornelas, & Press, 2000; Pickersgill, 2016). As far as we know, there are only two reports of tomato archeological remains, both seeds in coprolites: one in Southern Texas, close to the Mexican border, dated ∼2500 B.C. (Reinhard, Chaves, Jones, & Iñiguez, 2008), and one in the Peruvian Ica valley, from ∼500 A. D. In both cases the tomatoes were ingested, but there is no information regarding the degree of domestication (Beresford-Jones, Whaley, Ledesma, & Cadwallader, 2011).

Thus, despite past efforts, none of the tomato domestication models coherently captured all molecular, morphological, and passport data. To gain deeper insights into the domestication history of tomatoes, we developed a new approach based on an unsupervised automatic classification of haplotypic principal coordinate analyses (PCoAs) aligned via Procrustes (Krzanowski, 2000). The results of this analysis imply that SP evolved into SLC during a slow migration from Peru to Mesoamerica and that Peruvian and Ecuadorian SLCs are admixed populations originating from Mesoamerican SLC and Peruvian and Ecuadorian SP. Subsequently, Peruvian domesticated SLCs migrated north to Mexico and then evolved into SLL. Despite differing from the previously proposed evolutionary models, our model agrees with all available evidence, such as LD, genetic diversity, and distance, results from fastStructure (Raj et al., 2014) and TreeMix (Pickrell & Pritchard, 2012), and morphological and collection site ecological data. Moreover, other analyses have shown evidence that the selection of genes associated with photosynthesis and flowering time might have been involved in the long-distance migrations between different climatic types and latitudes.

## Materials and Methods

### Accessions and sequences

A total of 628 sequenced accessions, obtained from the Varitome project (Gao et al., 2019) and six previous studies (Causse et al., 2013; Lin et al., 2014; Sato et al., 2012; Strickler et al., 2015; Zhu et al., 2018) were included in the study. The sequences were obtained specifically from all SP, SLC, and SLL resequencing experiments publicly available in the Sol Genomics database (solgenomics.net) and the NCBI Sequence Read Archive, and three newly sequenced accessions of Ecuadorian origin. The passport data and mean coverage are listed in Tables S1 and S2.

### Mapping and single nucleotide polymorphisms (SNPs)

All reads were mapped with BWA-MEM (H. Li, 2013) against the latest available tomato reference genome (v.4.0) (Hosmani et al., 2019). After mapping, duplicate reads marked by Picard Tools (https://broadinstitute.github.io/picard/) and reads with a mapping quality lower than 57 were removed. For variant calling, the first and last three bases of every mapped read were ignored. SNP calling was conducted by FreeBayes, with a minimum coverage of 10, a minimum alternative allele count of two, an alternative allele frequency of 0.1, and a maximum number of searched alleles of five (Garrison & Marth, 2012).

After SNP calling, genotypes with coverage less than five were set to missing, and variants with a calling rate lower than 0.6, having observed heterozygosity greater than 0.1, or that were located on chromosome 0 were removed. Afterward, accessions with a calling rate less than 0.85 were removed from the analysis. Of the 25.3 million variants initially generated by FreeBayes, 11.8 million were retained after filtering out low-quality variants, and 2.02 million were present in the euchromatic regions. The variant filtering code is available in create_tier1_snps_excluding_low_qual_samples_and_chrom0.py.

Heterochromatic region determination was based on the recombination rate, and this rate was determined by interpolating data from the publicly available SolCAP genetic map constructed with 3,503 genotyped markers genotyped in the EXPEN 2000 F2 population (Sim et al., 2012). A region was considered heterochromatic when its physical distance-to-genetic distance ratio was lower than 1e-6.

We included all codes used to perform the analysis from variant filtering to figure creation in the supplementary materials and a public GitHub repository (https://github.com/bioinfcomav/tomato_haplotype_paper). The code developed was thoroughly tested, and the tests are also available in GitHub and can be run to check the correctness of the implementation. The tests for the calculated indices were also checked against standard population genetic software (Peakall & Smouse, 2012).

For most haplotypic analyses, the highly correlated heterochromatic regions were ignored, and to speed up the computations, only variants with at most 10% missing genotypes were used, resulting in a working set of 33,790 variants. For these haplotype analyses, the variants were phased and imputed using Beagle (Browning, Zhou, & Browning, 2018), and the relevant script used was phase_and_impute_with_beagle.py.

### Aligned haplotypic PCoAs and automatic haplotypic classification

The euchromatic regions were divided into segments. For each segment with at least 20 markers, a PCoA was conducted using the haplotypic alleles (two alleles per diploid individual) reconstructed after imputing and phasing. First, a pairwise distance matrix was constructed by calculating edit distances. PCoAs were then performed according to the methods of Krzanowski (Krzanowski, 2000). PCoAs from different genomic segments were then aligned using the SciPy orthogonal Procrustes function (Krzanowski, 2000). The Procrustes algorithm uses two sets of points in a space and then calculates and applies a linear transformation (e.g., rotates, translates, and/or reflects) to the second set of points to align it as best as possible to the first set. In this case, all the PCoA results were aligned to obtain the final set of aligned PCoA data. The code that implemented this functionality is located at haplo_auto_classification.py, haplo_pca.py, and procrustes.py.

Once all PCoA data were aligned, the haplotypes were automatically classified using an unsupervised classification algorithm. Before classification, the outlier haplotypes were removed using the isolation forest algorithm implemented by the scikit-learn Python library, with the contamination parameter set to 0.070 (Liu, Ting, & Zhou, 2012). Because of memory allocation limitations, outlier detection and unsupervised classification could not be conducted with the entire aligned PCoA haplotype matrix. This matrix has more than half a million rows, and these algorithms require a memory allocation that grows geometrically with the number of rows. To solve this problem, thinned input matrices were input into the algorithms. The thinned matrices were constructed by calculating the Euclidean distance between haplotypes, and when several haplotypes were closer than 0.0015, only one was retained in the thinned matrix. The automatic classification depended only on the haplotype location in the PCoA, such that once the classification was completed, all haplotypes that were close to the one present in the thinned matrix shared the same classification. We are aware that excessive thinning could alter the haplotype cluster density in the aligned PCoA data, which would affect the unsupervised classification. Therefore, care was taken to minimize the thinning. The thinning distance chosen was the minimum distance that created a matrix capable of being held in the computer’s memory. In total, 45,982 of the original 526,240 haplotypes were ultimately present in the thinned matrix.

Haplotypic unsupervised classification was performed in two steps. First, the haplotypes were classified using the agglomerative algorithm implemented by scikit-learn (Pedregosa et al., 2011). This agglomerative approach has one limitation: it forces the classification of each haplotype. To solve this problem, the automatic classification was refined in a second step by the KNeighbors supervised classification algorithm that uses the classification generated by the agglomerative step as an input. KNeighbors was configured to use 30 neighbors (Pedregosa et al., 2011).

### Population structure

The accessions were classified into populations accounting for their taxonomic and collection origin passport data and a series of variant-based (not haplotypic-based) hierarchical PCoAs. A population was defined when a set of contiguous accessions in a PCoA were collected from the same geographic area or were classified in the same taxon and shared a similar haplotype composition inferred from the previously calculated haplotypic-based PCoAs. The variant-based PCoAs were performed only with variants with a major allele frequency lower than 0.95, a missing genotype rate lower than 0.1, and presence in the euchromatin. To avoid overrepresentation of any genomic region, the euchromatin was divided into segments of 100 kb, and from each segment, only the most variable variants were retained. Using the resulting variants, the Kosman distances were calculated (Kosman & Leonard, 2005), and a PCoA was conducted (Krzanowski, 2000). The relevant scripts used include both pcas_do.py and pca.py.

FastStructure was also used to infer population composition (Raj et al., 2014). This method was run with Ks ranging from two to 11 with the default settings. For this analysis, only the variants with a maximum major allele frequency of 0.95 and were located in the euchromatic regions were used. The code used was embedded in the fastructure_run.py.

### Parametric population history reconstruction

We used TreeMix to reconstruct the population history using variants characterized by low LD (Pickrell & Pritchard, 2012). Thus, the variants were selected by generating haplotype blocks of consecutive SNPs in the genome that allowed only a maximum correlation threshold between its genotypes. From every haplotype block, only one variant was selected at random for use by TreeMix. Branch support was calculated by bootstrapping, and in each bootstrap iteration, random variants were used from every linked genomic block. Different TreeMix analyses with different LD thresholds were performed to test the robustness of the results. The code can be found in the treemix.py file.

### Population diversity and LD

To evaluate the population diversity, various parameters were calculated, such as the number of polymorphic variants (95% threshold) and unbiased Nei diversity (Nei & Roychoudhury, 1974). Additionally, to reflect the haplotypic diversity, several indices were evaluated for each 500 kb genome segment: the mean number of different haplotypes or the mean number of variants found in a genomic segment.

Some of these parameters could potentially be affected by the number of individuals available in each population. Two strategies accounted for this potential pitfall. In the first strategy, diversities were calculated using the same number of accessions for each population. The number of accessions was chosen to be 75% of the number of accessions of the population with the fewest accessions in the analysis. The indexes were then calculated 100 times, with different accessions randomly chosen for each population in each iteration. When all 100 values were obtained, the mean and confidence interval of the mean were used to represent the diversity of each population. In the second approach, a complementary rarefaction analysis, in which accessions were added one by one to each population, was performed. The script files used to perform these calculations were diversities_vars.py and diversities_haplos.py, respectively.

The LD was calculated using only polymorphic variants (95% threshold). LDs between markers that were up to 1000 kb apart were calculated, and LD decay was estimated using locally weighted scatterplot smoothing (LOWESS) implemented by the statsmodels Python library (www.statsmodels.org). Finally, the LD of 10 kb was obtained from the adjusted curve. The relevant code is located in ld.py, ld_decay_plot.py, and ld_bar_plot.py.

### Introgression detection and functional analysis

ABBA-BABA indices were calculated using euchromatic variants (Green et al., 2010). Additionally, the mean values per genome region were obtained and plotted to look for introgressions in Ecuadorian and Peruvian SLC. These genomic regions were 50 kb in length. The relevant scripts are abba.py and there_and_back_abba.py.

Additionally, the alleles potentially introgressed from SP into SL were examined using the method below. Alleles in the SLC MA population were considered reference SL alleles, and alleles found in all other SLC populations were labeled as introgressed when they were not found in SLC MA but were present in an SP population at a frequency higher than 10%. SLC MA was used as a reference because the results showed that this population had the fewest Peruvian and Ecuadorian haplotypes. Once each allele for each variant was labeled as introgressed or not introgressed, the allele introgression frequency was calculated for each variant. The implementation can be found in the introgressions.py file.

### Morphological analysis

Morphological analysis based on a characterization of 375 accessions available at the Tomato Genetic Resource Center (TGRC) and the COMAV GenBank was performed using the accessions for which images were available. Each accession was evaluated for basic inflorescence, leaf, stem, and fruit traits (Table S1, Fig. S15).

Morphological classification was conducted via principal component analysis (PCA). Morphological traits were treated as ordinal traits. For the PCA, accessions with more than seven missing values were removed, and any missing data in the remaining accessions were filled by the means. The data for each trait were standardized using the StandardScaler function of the scikit-learn Python library, and PCA was performed using the PCA functionality implemented in the same library (Pedregosa et al., 2011). The code is located in the morphological.py file. Finally, morphological classification was performed manually by inspecting the morphological PCA data and the taxonomic and collection site passport information.

The passport data were manually curated to determine the collection source information. The final collection source for each accession was obtained by combining the information stated in the collection source passport field and, when available, the annotations and images made during the collection expeditions.

## Results

### Haplotypic PCoAs and automatic classification

The euchromatic regions of the entire genome were divided into 440 segments (0.5 Mb), resulting in a total of 526,240 haplotypes (two per plant accession per 440 segments). The output of the PCoAs conducted with the haplotypes of each segment was aligned using Procrustes, resulting in a triangular-like structure that matched the structure of the main taxonomic groups (Fig. 1). The haplotypes were automatically classified into three types using an unsupervised clustering algorithm. The three haplotype types were named according to the taxonomic groups in which they were usually found: hPe (most abundant in Peruvian SP), hEc (most abundant in Ecuadorian SP and SLC), and hSL (most commonly found in SLC and SLL). The chosen number of haplotype types, three, was in good agreement with the number of ancestral populations suggested by the fastStructure marginal likelihoods (Fig. S2). However, the Calinski-Harabasz (CH) clustering score, an index commonly used to evaluate clustering performance, suggested fewer types (Fig. S2). This might have been caused by the intermediate haplotypes found between the main clusters, likely caused by recombinant and intermediate haplotypes and by the uneven representation of the different haplotype types. SLL accessions were overrepresented, with 44% of the sequenced accessions, whereas the SP Ec, a traditionally ignored population, was underrepresented with only 2.5% of the accessions. To determine the effect of the number of haplotype types, haplotype classification was also performed with two, three, four, and five haplotype types (Fig. S3). When two types were used, the haplotype mainly divided SP from SL, whereas when more types were allowed, SP was divided into subtypes, such as Ecuadorian and Peruvian SP for three types. The relationship between SL and SP, the main focus of the current analysis, remained unchanged when the number of haplotype types was altered.

**Fig. 1.**
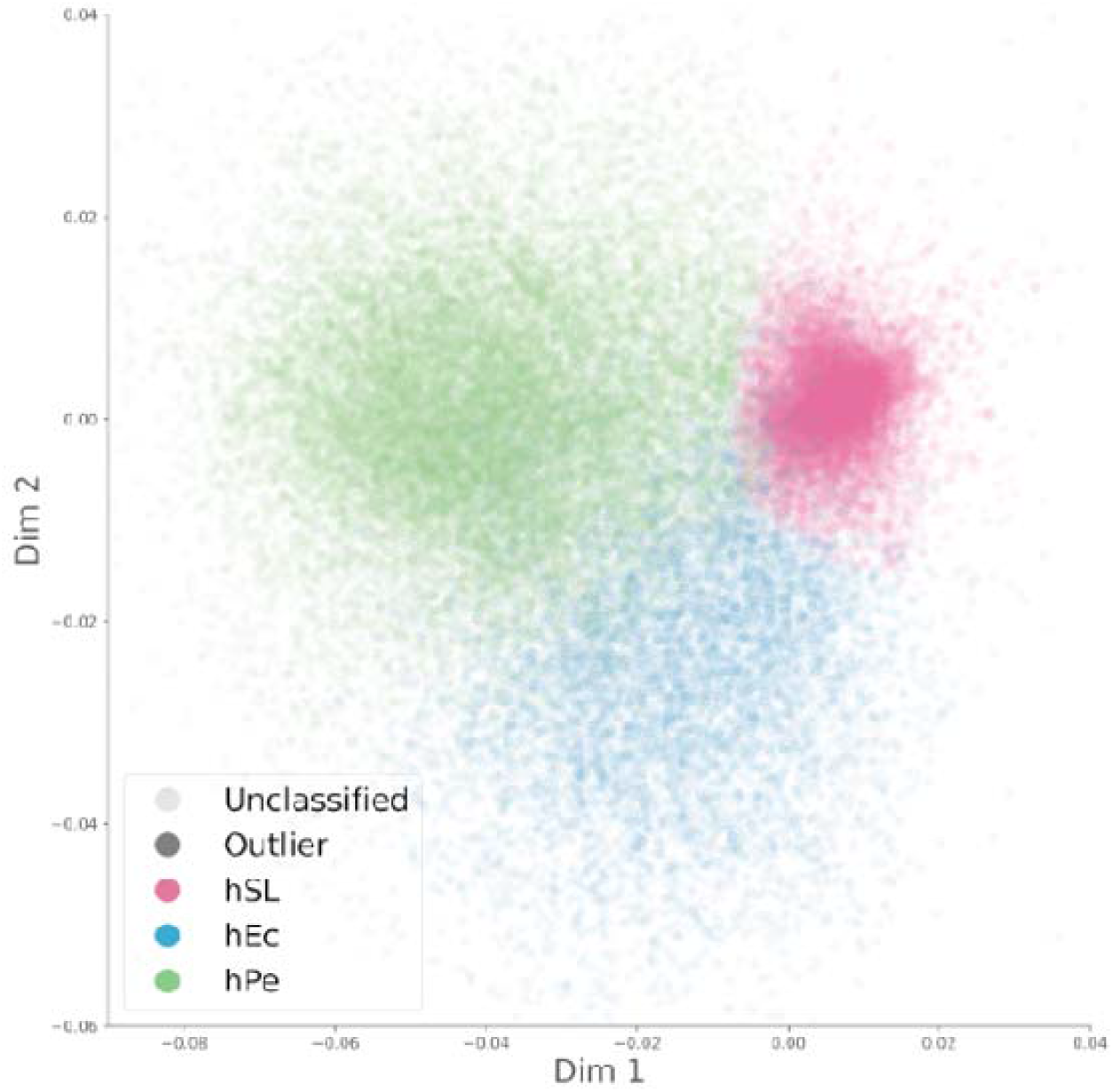
Haplotypic PCoAs. A PCoA was conducted for every 500 kb genome segment using edit distances between haplotypes. The resulting PCoAs were aligned using Procrustes and automatically classified into three haplotype types.

Haplotypic classification was used to analyze the genomic composition of each accession (Fig. S4 and Fig. S5). For example, the haplotype composition for the Cervil accession (Causse et al., 2013) is shown in Fig. S6. The haplotype classification and location in the aligned PCoA results showed that this accession comprised hPe and hSL haplotypes. Therefore, it is likely that Cervil is the result of hybridization between Peruvian SP and cultivated tomato.

### Accession classification and haplotype composition

The accessions were classified into genetic groups through a series of hierarchical PCoAs calculated from the genetic distance matrix (Fig. 2 and Fig. S7), and the information provided by the geographic and taxonomic passport data (Tables S1 and S2) and their haplotype composition (Figs. S3, S4, and S5). SP was split into three populations: SP Pe (Peru), SP Montane, and SP Ec (Ecuador). The most abundant SLC populations were SLC MA (Mesoamerica), SLC Pe, and SLC Ec. SLL was composed of SLL Mx (Mexico), SLL vintage, and SLL modern (Table S2). Other minor populations were noted, such as SLC Co (Colombia), but they were represented by only a few accessions; thus, they could not be used in all analyses. The overall genetic group separation was similar to that used in previous studies (Fig. S8) (Razifard et al., 2020).

**Fig. 2.**
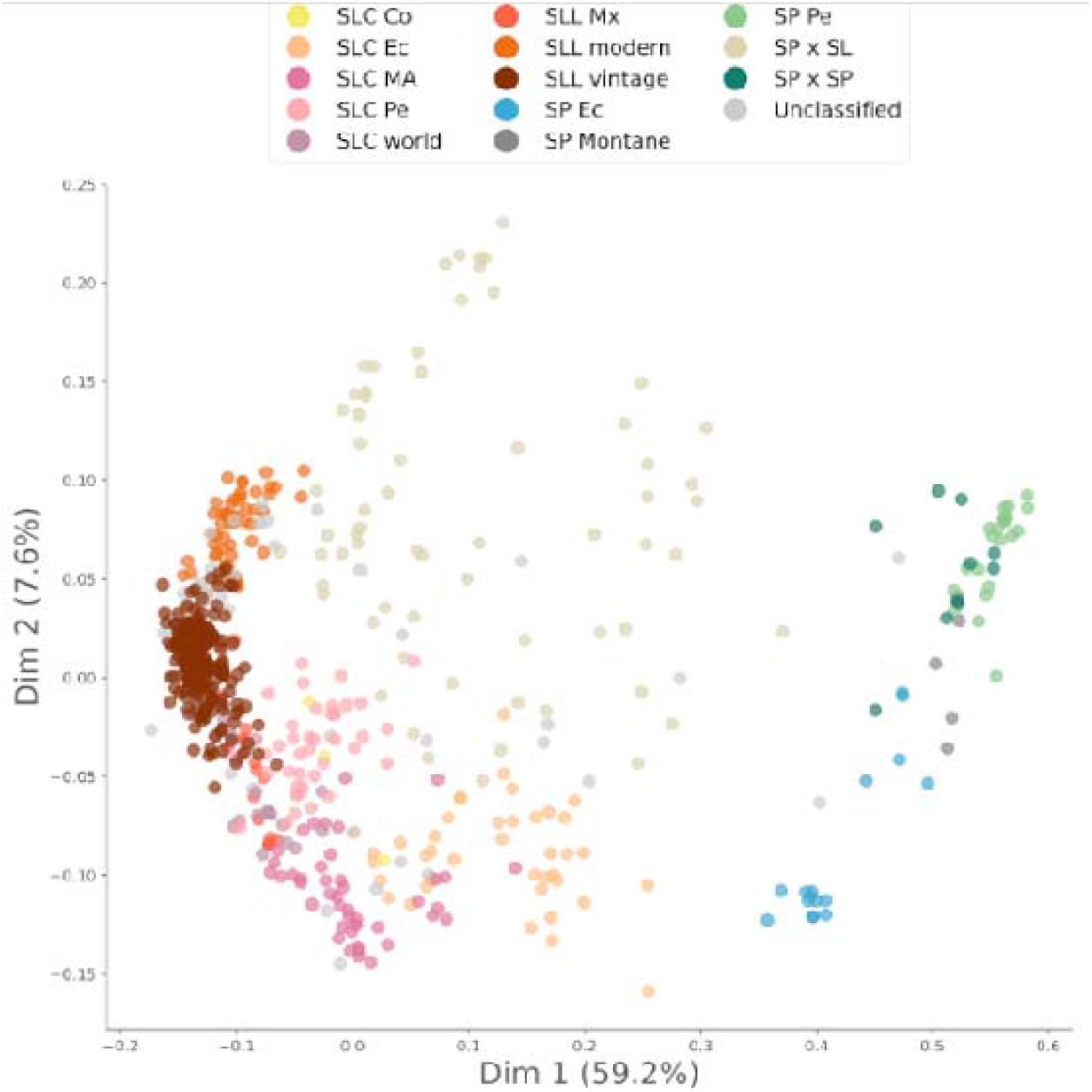
Accession PCoAs. Pairwise Kosman genetic distances between accessions were calculated using variants characterized by a major allele frequency lower than 0.95 and a missing genotype rate lower than 0.1. Only the most variable variant from every 100 kb genomic segment was used to calculate the distances. The PCoA was based on the obtained pairwise genetic distance matrix. The accessions are colored according to the population in which they were classified.

The haplotype composition of each population was assessed by plotting the haplotypes of the accessions belonging to each population in the aligned PCoA (Fig. 3 A). The ancestral population composition calculated from the fastStructure results was similar to the haplotype composition at the population (Fig. 3 B and C) and individual level (Fig. S5).

**Fig. 3.**
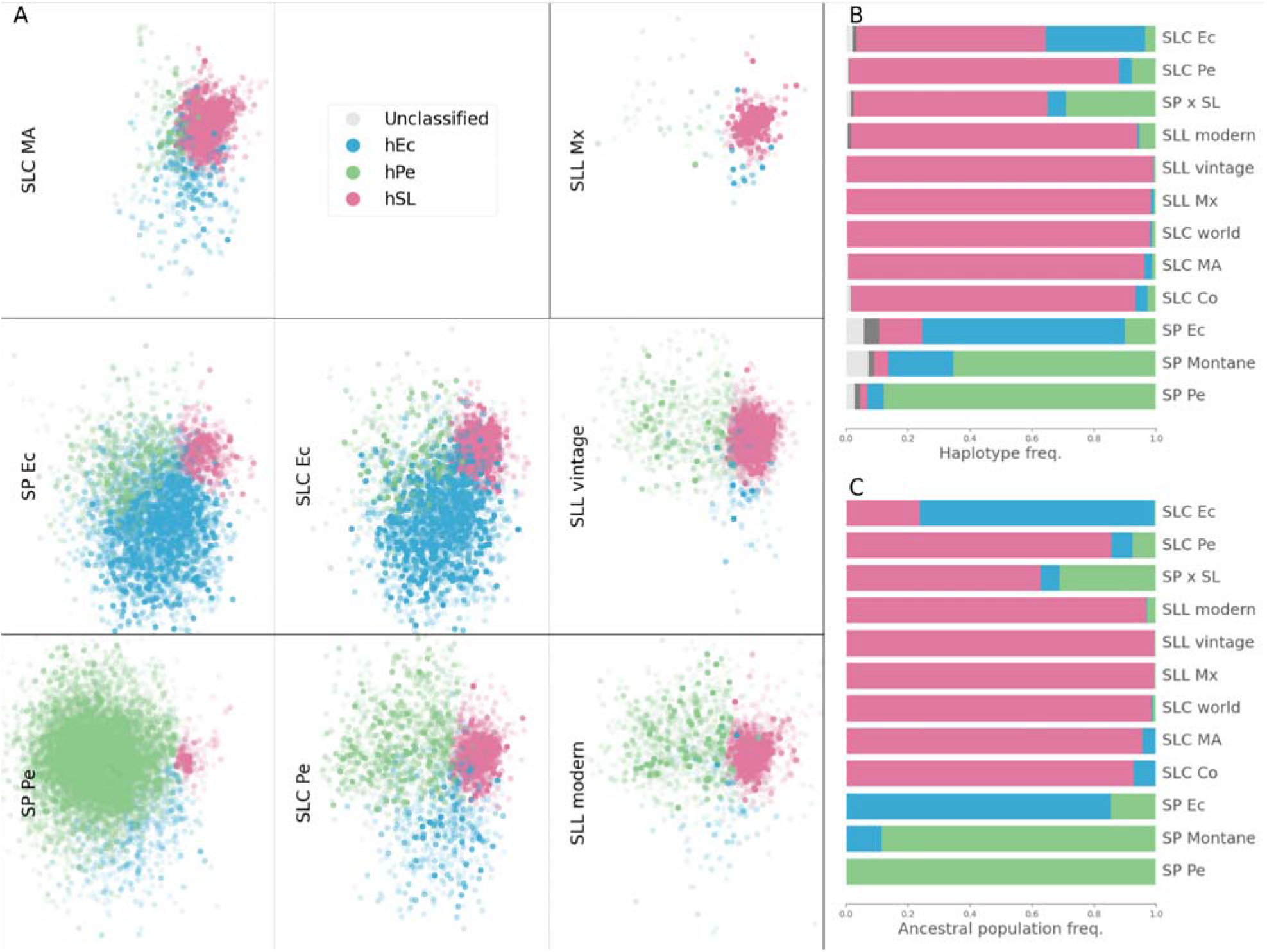
Population haplotype composition. A) Classified haplotype PCoA obtained for Figure 1 was divided into several figures, one per population. In each figure, only the haplotypes that belong to accessions for each population were included. B) Frequencies of each haplotype type found in each population. C) FastStructure ancestral population composition per population.

To determine if the size chosen for the genome segments could affect the haplotype analyses, they were repeated with different genome segment sizes (100, 500, and 1000 kb), and the results were similar in all cases (Fig. S9).

The genetic diversity of the haplotypes that belong to one of the haplotype types (hPe, hEc, hSL) was calculated for each population (Fig. 4). SP Pe was the most diverse population for haplotype type hPe, SP Ec for hEc, and Mesoamerican SLC for hSL. These diversity results agreed with the fastStructure results, which also suggested that three haplotype types (hPe, hEc, hSL) corresponded to the three ancestral populations (Fig. 3 C). Furthermore, the haplotypes associated with hPe and hEc were most abundant in the extant SP Pe and SP Ec populations, respectively (Fig. 3 B). The SL populations were genetically close and contained mostly hSL haplotypes.

**Fig. 4.**
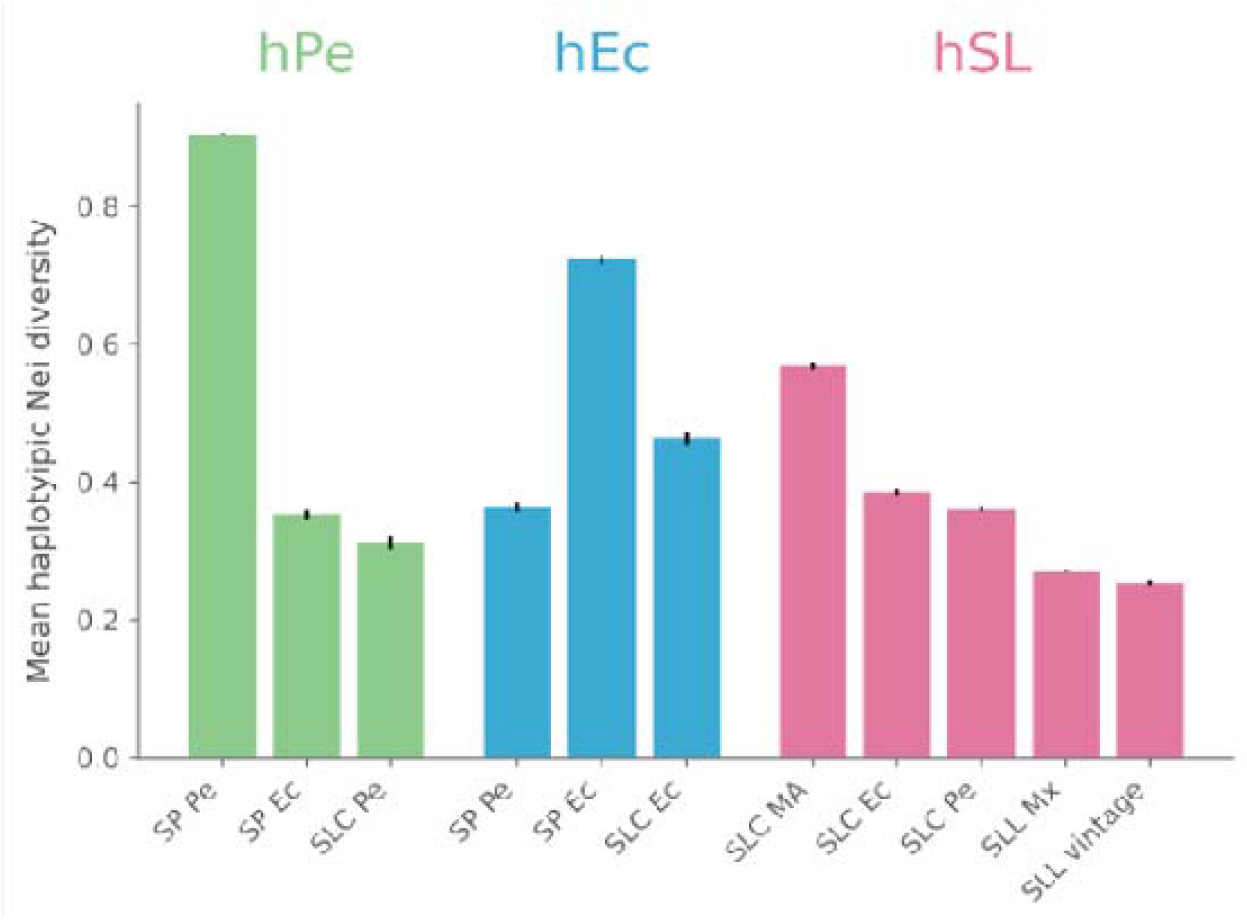
Mean Nei haplotypic diversity per haplotype type. Euchromatic regions were split into 500 kb segments; for each segment, the haplotypic alleles were determined and classified. The unbiased expected heterozygosity per variant was calculated using only the genotypes corresponding to the haplotypes classified as hPe, hPEc, and hSL. The mean of the expected heterozygosity for the whole genome was calculated. To calculate the indexes, the number of accessions was the same for each population, being 75% of the population with fewer individuals. The analysis was repeated 100 times, choosing the accessions representative of each population at random. The bar represents the mean obtained in the 100 repeats, and the error bars are the confidence intervals of the means.

Overlap of the divisions among Peru, Ecuador, and SL, the population haplotype composition had an evident pattern of secondary contacts. For instance, although most haplotypes found in SLC Pe were hSL, certain hPe and hEc haplotypes were also present in this population. Remarkably, a quarter of the haplotypes found in the Ecuadorian SLC were not hSL but hEc haplotypes; thus, they might have been introgressed from SP Ec. To further investigate the complex patterns of gene flow, we employed TreeMix (Fig. S10). According to the TreeMix analysis (Fig. S10 A) SLC MA appeared to be basal to all other SLCs. SLC Pe and SLC Ec were then derived from SLC MA by acquiring introgressions from SP Pe and SP Ec, respectively. These results were also in agreement with the ABBA-BABA analyses (Green et al., 2010). For SLC Ec, SLC MA, SP Ec, and SP Pe, the D statistic was -0.23, whereas for SLC Pe, SLC MA, SP Ec, and SP Pe, the D value was 0.43. In both cases, the p-value was 0. These D statistics were compatible with SLC Ec receiving introgressions primarily from SP Ec, whereas SLC Pe would have introgressed genomic segments mainly from SP Pe.

### Diversity and LD

The overall number of polymorphic (95% criteria) genetic variants (Fig. 5 A), the mean number of variants found in a genomic region (Fig. 5 B), and the unbiased Nei genetic diversity (Fig. S11) yielded similar results. According to these indexes, the most diverse populations were the Peruvian and Ecuadorian SP and SLC and modern SLL. A relatively low genetic diversity characterized SLC MA, SLC World, and particularly SLL Mx. These results matched those of previous analyses (Jose Blanca et al., 2012; José Blanca et al., 2015), but contrasted with those of another diversity measure, the mean number of different haplotypes found in a given genomic region (Fig. 5 D). According to this index, the SP populations were the most diverse, whereas all SLC populations exhibited lower levels of diversity. Although SLC Ec and SLC Pe were seemingly highly variable according to the number of polymorphic variants, these populations did not have many different haplotypes. These results also indicated that SLC Ec and SLC Pe might result from an admixture of an ancient SLC population with an SP, as already suggested by the haplotype composition, the fastStructure, the TreeMix, and the ABBA-BABA analyses. Furthermore, LDs of recently created populations or populations that recently incorporated genetic material were usually high, and the SLC population with the lowest LD was SLC MA (Fig. 5 C).

**Fig. 5.**
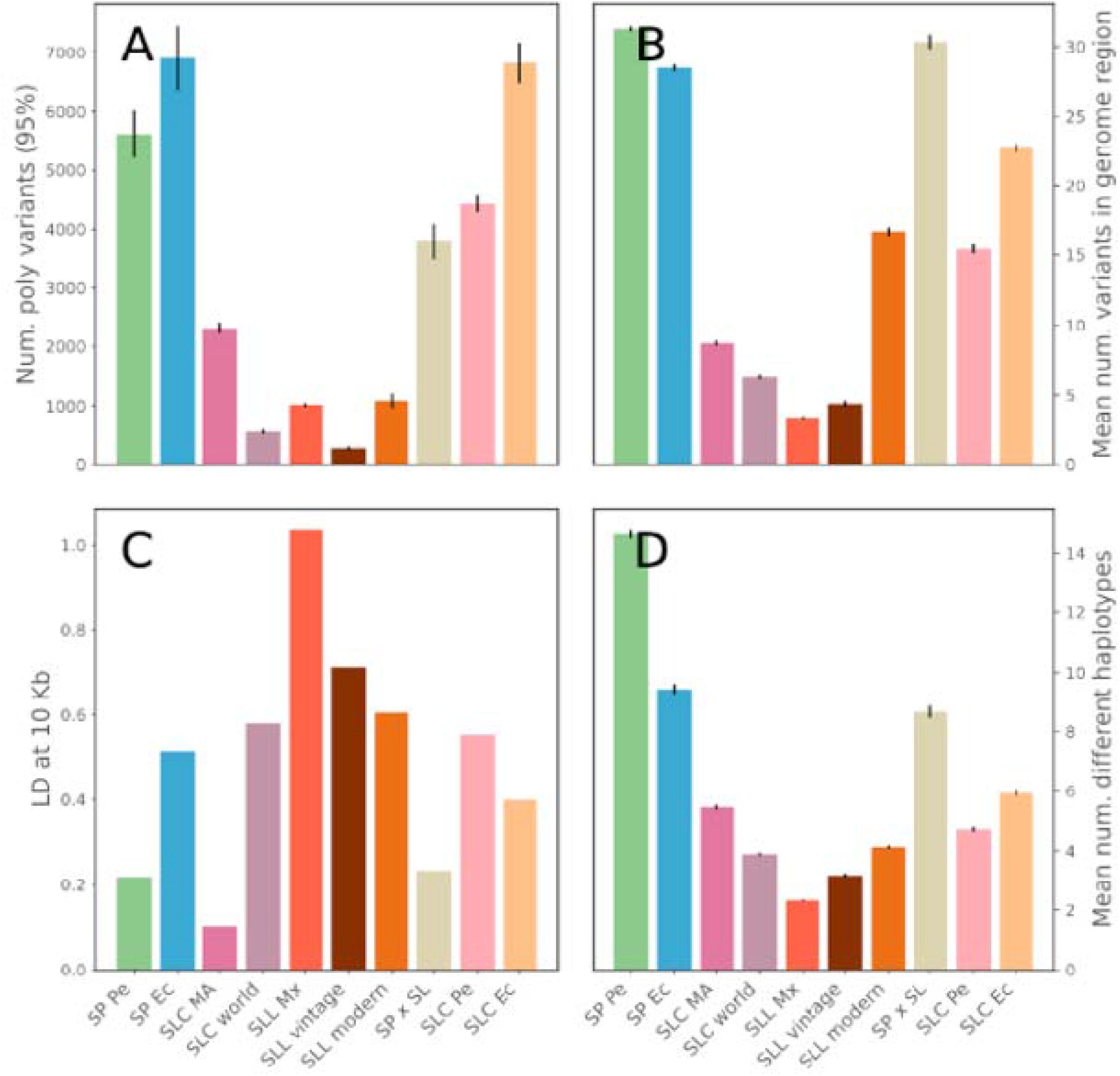
Diversity and linkage disequilibrium per population. To calculate the indexes, the number of accessions was the same for each population, 75% of the population with fewer individuals. The analysis was repeated 100 times, choosing the accessions representative of each population at random. The bar represents the mean from the 100 repeats, and the error bars are the confidence intervals of the means. A) Number of polymorphic variants (95% threshold). B) Mean number of variants found in a 500 kb euchromatic segment. C) Linkage disequilibrium between variants at 10 kb. D) Mean number of haplotypic alleles (500 kb euchromatic segments).

### Latitude-related selection

According to all previous researchers and all the evidence presented, SP Pe is the oldest population; thus, SLC MA would result from northward migration. In this migration, some genomic regions could have been selected. To study the possibility of a selective sweep, the expected heterozygosity was calculated along the genome for SLC MA (Fig. 6 A). SLC Ec and SLC Pe appeared to be derived from SLC MA; however, according to the haplotype, TreeMix, fastStructure, and ABBA-BABA analyses, both have introgressions from SP. Thus, we calculated the D, BABA, and ABAA indices along the genome assuming the following evolutionary schemas: 1) SLC Ec, SLC MA, SP Ec, and SP Pe and 2) SLC Pe, SLC MA, SP Ec, and SP Pe (Fig. 6 B, Fig. S12). Some genomic regions have an introgression frequency, such as the chromosome 1 end, or the region just before the pericentromeric region on chromosome 4. We analyzed the possible relationship between the diversity in SLC MA and the introgression rate in SLC Ec (Fig. 6 C). The regions more introgressed in SLC Ec had lower diversity in SLC MA. This result appeared to indicate that the selection process experienced during the northward migration might have been partially reversed by introgressions from SP after the Ecuadorian colonization by SLC. Additionally, we tested this hypothesis by calculating the frequency of SP alleles in SLC Ec that were not present in SLC MA, and we related these possible introgressions with the SLC MA diversity (Fig. S13). In this analysis, the genomic regions with an abundance of SP alleles were also correlated with regions with lower diversity in Mesoamerican SLCs.

**Fig. 6.**
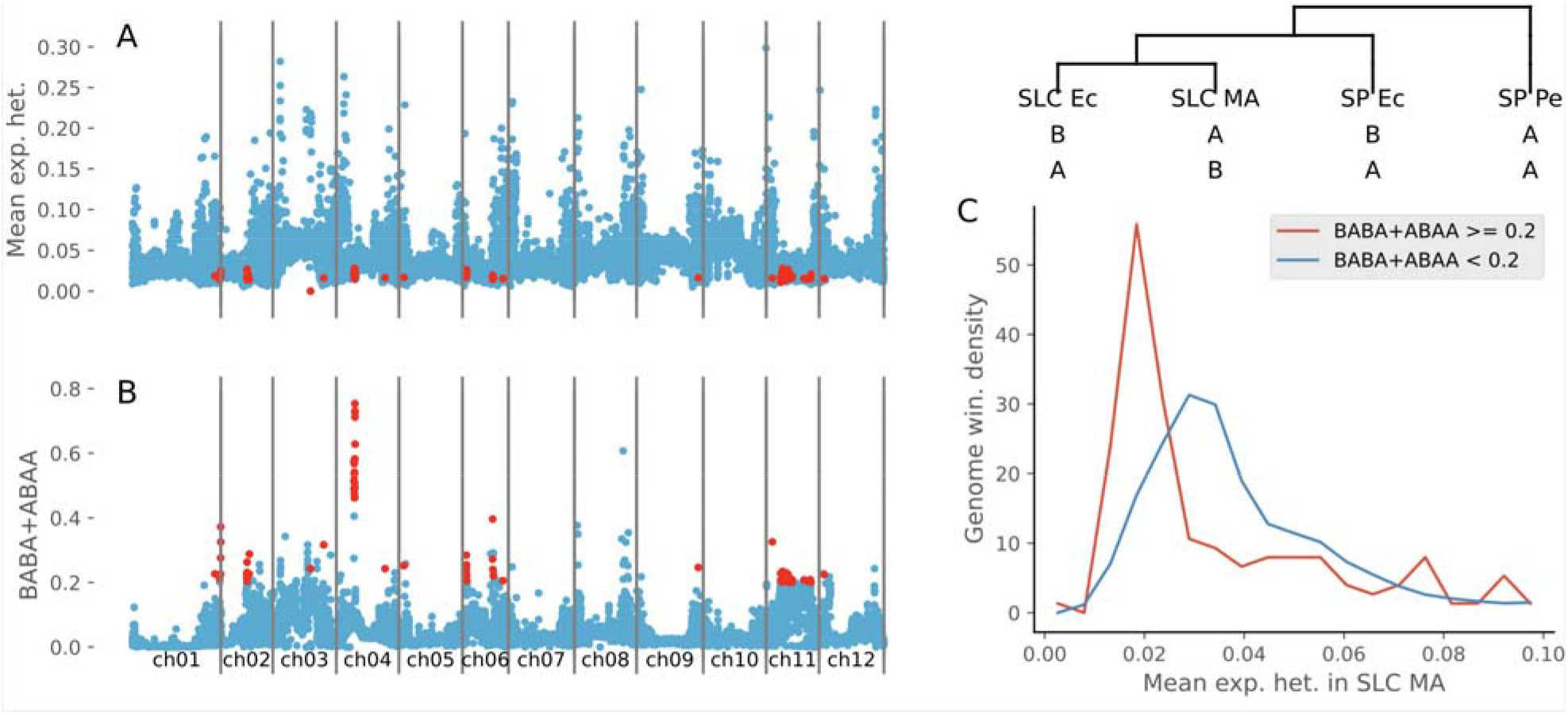
Mesoamerican Nei diversity and introgression detection in SLC Ec along the genome. A) Mesoamerican SLC expected heterozygosity along the genome in 500 kb segments. B) Sum of BABA and ABAA products calculated in 500 kb segments along the genome using the evolution model: SLC Ec, SLC MA, SP Ec, SP Pe. The genome segments with a BABA + ABAA value higher than 0.2 and expected heterozygosity lower than 0.03 are represented in red. C) SLC MA expected heterozygosity genome segment distributions with BABA + ABAA lower and higher than 0.2.

We inspected the genes found in regions with high introgression rates in SLC Ec and SLC Pe and low diversity in SLC MA (Table S3). Some regions, such as those on chromosomes 2, 8, and 11, were large and comprised hundreds of genes, whereas others, such as those on chromosome 7, were smaller. Only five genes were identified in this region, one of which was Solyc07g043270, a FAR-red elongated hypocotyl 3-like protein-encoding gene. FAR-red genes respond to light and have been related to flowering time and other processes regulated by light conditions (G. Li et al., 2011; Xie et al., 2020). FAR-red genes were detected in three of these regions. On chromosome 4, two of the 22 genes found were a spermidine synthase (Solyc04g026030) that might also be involved in the regulation of flowering time (Imamura, Fujita, Tasaki, Higuchi, & Takahashi, 2015). Additionally, an Agamous protein was possibly involved in flowering and fruit development (Pan, McQuinn, Giovannoni, & Irish, 2010). In total, three Agamous genes were found in these regions. On chromosome 2, Solyc02g021650, a component of the light signal transduction machinery involved in the repression of photomorphogenesis (Lieberman, Segev, Gilboa, Lalazar, & Levin, 2004), and on chromosome 6, Solyc06g050620, a reticulata-related family gene associated with chloroplast development, among other processes, was detected (Pérez-Pérez et al., 2013).

### Mexican SLL origin

To further determine the relationship between the Mesoamerican and Peruvian SLCs and the Mexican SLL, a detailed haplotype-based analysis was conducted (Fig. 7). The populations with the most private haplotypes were the Mesoamerican SLC and Peruvian SLC populations. Many private Peruvian SLC haplotypes appeared to be the result of introgressions from Peruvian and Ecuadorian SP. The Mexican SLL had the fewest private haplotypes. The two population pairs that shared the most haplotypes were Mesoamerican and Peruvian SLC and Peruvian SLC and Mexican SLL; however, despite their geographic closeness, SLC MA and SLL Mx shared fewer haplotypes. Thus, according to these results, it is plausible that Mexican SLL originated from the Peruvian SLC and not from the geographically closer Mesoamerican SLC.

**Fig. 7.**
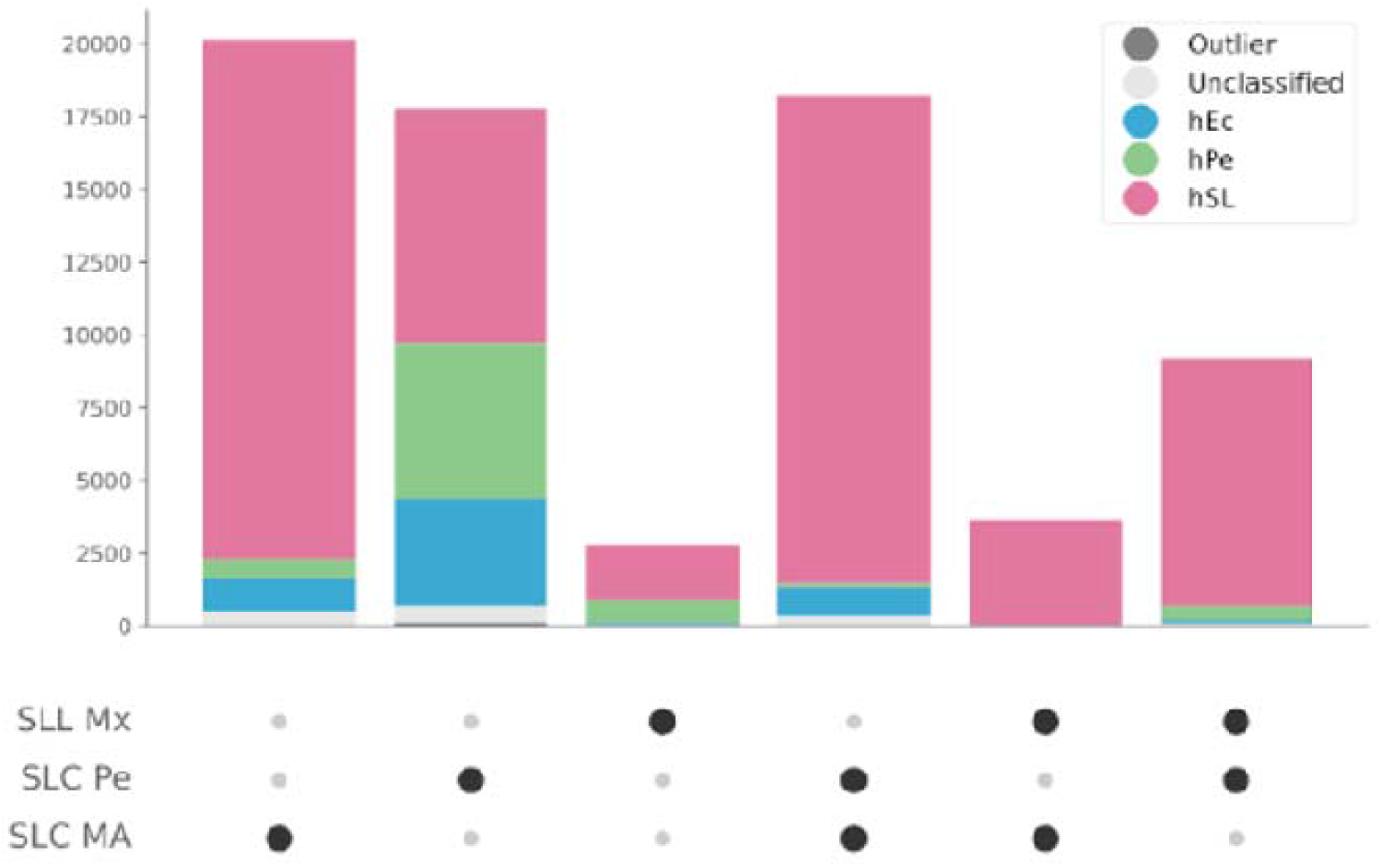
Distribution of haplotypic alleles. Shared and private haplotypic alleles among Mesoamerican SLC, Peruvian SLC, and Mexican SLL. The bars represent the number of the different haplotypes shared between the populations, with higher numbers depicted by the larger dots in the lower panel.

### Morphological analysis

Morphological characterization of SP, SLC, and SLL was also conducted. A PCA for leaf-, inflorescence-, fruit-, and stem-related traits was used to cluster accessions into several morphological types (Fig. 8 A, Fig. S1, Fig. S14, and Fig. S15). Three morphological types were found in SP, of which the first featured longer inflorescences, wider petals, and frequently, striped fruits, and curved and exerted styles compared with those of the other types. This first morphological type was characteristic of the Peruvian SP. Compared with the other types, the second type comprised mostly northern Ecuadorian SP accessions and featured slightly larger fruits, which usually showed a distinct transversally elongated shape (peanut-like shape). Finally, the intermediate SP morphological type found in Peru and Ecuador was characteristic of accessions found in the mountainous valleys between Peru and Ecuador. SLC was mainly divided into three morphological types: SLC small, SLC big, and SLC Ecu. SLC small had small fruits, with sizes ranging from small SP fruit size to cherry size, whereas SLC big had relatively large fruits that varied in size from cherry size to full-size commercial fruits. These relatively large fruits were frequently ribbed and almost always flat. Fruit size was found to be associated with stem width. Finally, the endemic Ecuadorian SLC form was characterized as having a mixture of characteristics between different SLC and SP types: it had cherry-sized fruits like those of SLC but folded back petals and longer inflorescences, similar to the intermediate SP type, and no stem pilosity and some transversally elongated fruits, such as those of Ecuadorian SP.

**Fig. 8.**
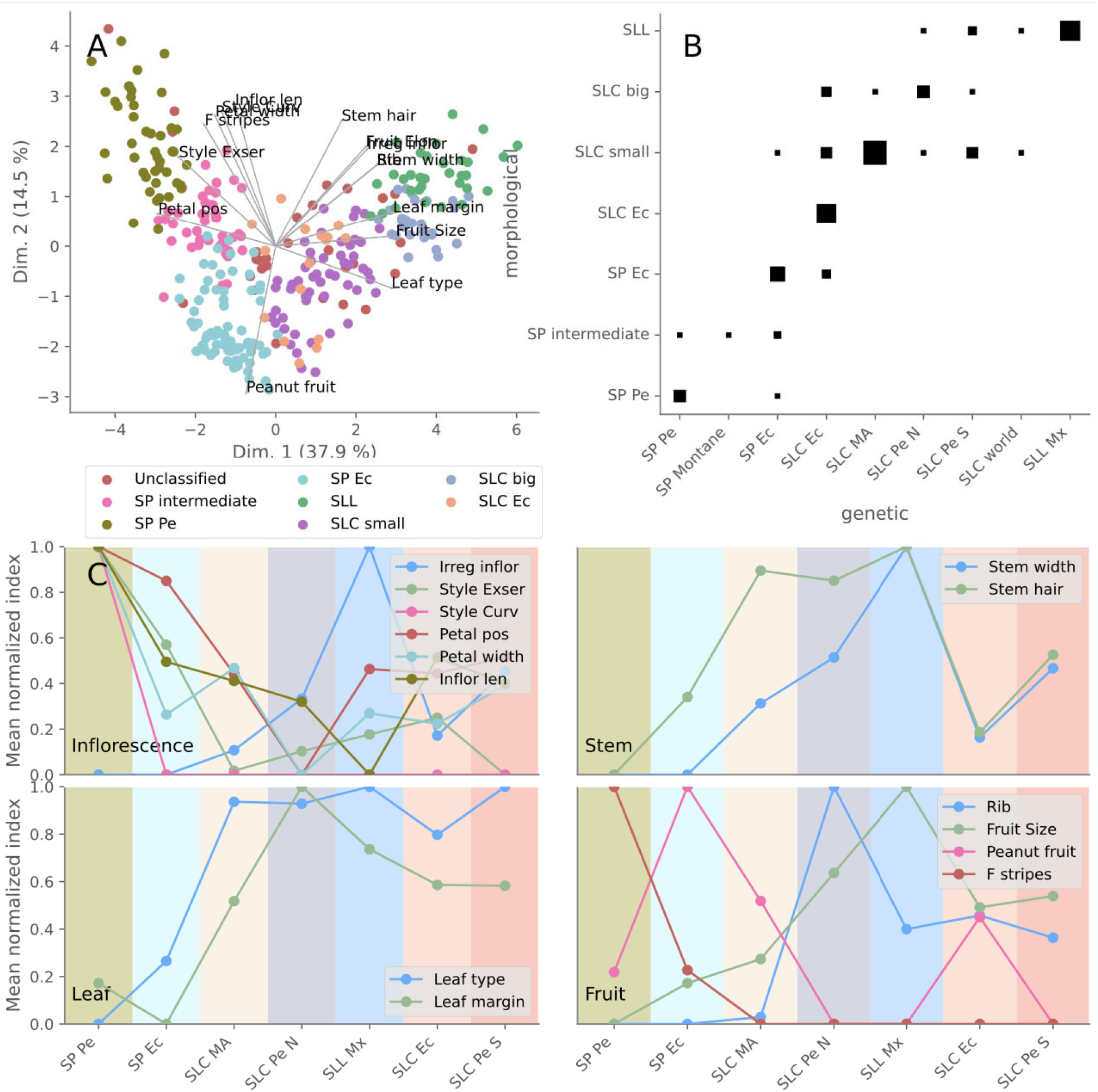
Morphological analysis. A) Accession-based Principal Component Analysis calculated using the morphological traits. The marker colors represent different morphological types. B) Comparisons between morphological types and populations. The marker size represents the number of accessions. C) Morphological trait mean values calculated for different populations.

These morphological types were associated with molecularly defined populations (Fig. 8 B). For this analysis, SLC Pe was divided into two genetic subpopulations, northern Peruvian SLC (SLC Pe N) and southern Peruvian SLC (SLC Pe S) (Fig. S7 B and C), which exhibited differential morphological characteristics, despite being closely genetically related. The SP Pe and SP Ec genetic groups roughly corresponded to the Peruvian and Ecuadorian SP morphological types. The SP Montane population was characterized by the intermediate type, although this type was also found in other SP populations. Mesoamerican SLC primarily produced small fruits, whereas northern Peruvian SLC was characterized mainly by SLC big or SLL morphological types. The other American SLC populations, southern Peruvian SLC and Ecuadorian SLC were morphologically variable and included morphological types that produced large and small fruits, and in the Ecuadorian case, also the typical Ecuadorian SLC-type.

Additionally, an analysis of the collection sites taken from the passport data was performed (Fig. S16). Undisturbed environments were labeled natural, whereas accessions collected in human-altered environments, such as roadsides, were considered ruderal. Semi-cultivated or cultivated accessions were collected mainly from backyards. The collected data were manually curated, and sometimes, these passport data were complemented by the information obtained from images of the collection sites. A total of 262 accessions had associated collection data. Most SP accessions (160 vs. 10) and 65% (17 vs. 9) of the SLC MA accessions were wild or ruderal, whereas the rest were weedy or semi-cultivated. These findings contrasted with the data concerning Ecuadorian SLC, which was semi-cultivated or cultivated in 89% of the occasions (25 vs. 3 accessions).

## Discussion

Previous studies have reported a complex history of SLC, the wild and semi-domesticated variety related to the cultivated tomato (Fig. 9) (Jose Blanca et al., 2012; José Blanca et al., 2015; Razifard et al., 2020; C. M. Rick & Holle, 1990). However, even with the available genomic evidence, no detailed tomato evolutionary model capable of accounting for all the empirical evidence has been produced. In the current analysis, the wild and cultivated genetic diversity present among SP, SLC, and SLL were analyzed considering all the publicly available whole-genome resequencing data mapped to the latest tomato genome reference (v4.0) (Hosmani et al., 2019), as well as the morphological and passport data gathered from various gene banks. Moreover, we developed a novel method to perform a genome-wide haplotypic analysis by combining Procrustes-aligned PCoA output with automatic unsupervised classification. This new method allowed a detailed and quantitative inspection of the haplotype composition of each accession and population; thus, it was useful for studying gene flow, introgression, and migration without the need for any assumptions related to the Hardy-Weinberg equilibrium or reproductive system of the species involved. The only limitation was sufficient LD; otherwise, the haplotypes would not be informative. To our knowledge, Procrustes has never been used for this purpose, and its use in population genetics has been restricted to the alignment of genomic PCA data and geographic maps (Wang et al., 2010) or alignment of PCA data generated from different SNP datasets (Wang, Zhan, Liang, Abecasis, & Lin, 2015). The domestication model obtained was supported by traditional population genetic indices, parametric statistical models, and morphological and passport data. We hope that similar approaches can be used to study the complex domestication histories of other species.

**Fig. 9.**
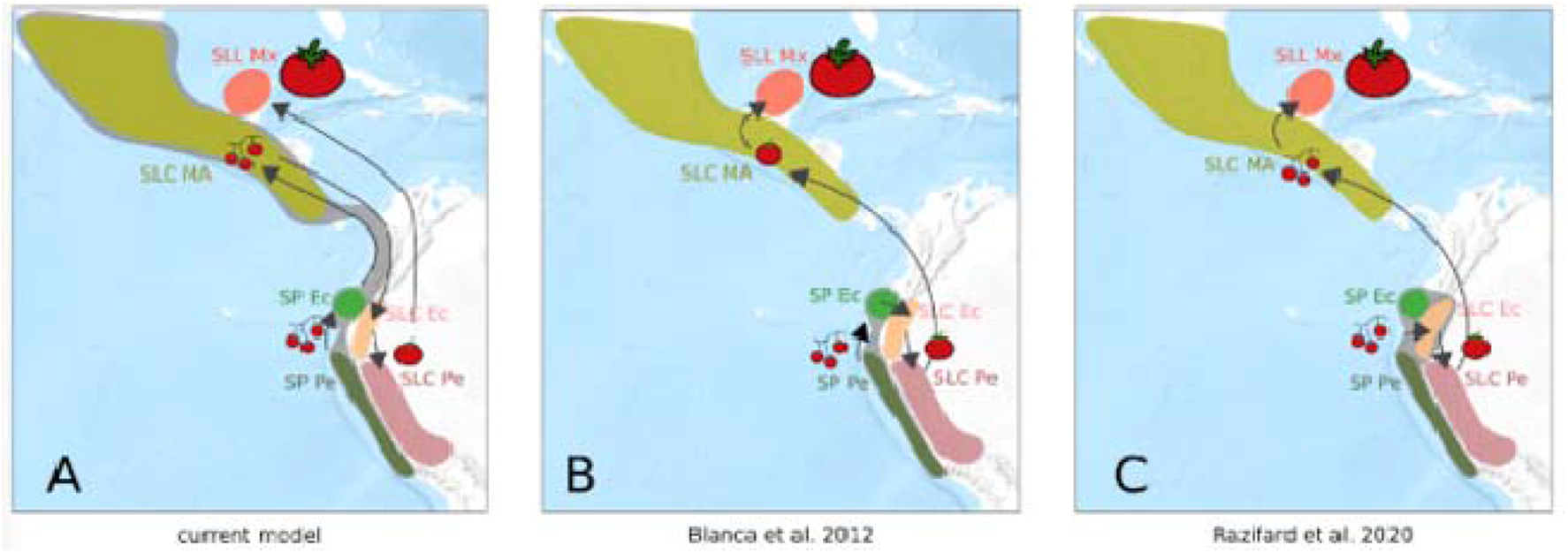
Tomato evolution hypotheses. Genetic population ranges are represented by the coloring of the different geographical areas. The areas in grey include populations that evolved before any human alteration. Arrows indicate migrations. Three fruit sizes are represented: wild-like small fruits, semi-domesticated forms, and SLL fruit types. A) Evolutionary model proposed in the current study. B) Hypothesis proposed by Blanca et al. 2012. C) Hypothesis proposed by Razifard et al. 2020.

### Three original wild populations

This novel analysis provided empirical evidence that suggests a tomato evolutionary model that accounts for all previous problematic results (Fig. 9 A). The haplotypic PCoA results, in agreement with fastStructure, suggested the existence of three haplotype types (hPe, hEc, and hSL) related to the main tomato taxonomic groups (Fig. 3 B and C and Fig. S5). The haplotype composition, fastStructure, and genetic diversity of each type of haplotype (Fig. 4) suggested that the three haplotype types (hPe, hEc, and hSL) originated in three ancient populations related to the extant SP Pe, SP Ec, and SL MA populations (Fig 9. A). SP Pe, the only population, composed mainly of hPe haplotypes, was also the most diverse population. Owing to its diversity and abundance, this population has been considered in all previous studies as the center of origin for SP (Charles M. Rick & Fobes, 1975). hEc haplotypes were mainly present in the two Ecuadorian populations, namely, SP Ec and SLC Ec (Fig. 3 A and B), but they were more common and diverse in SP Ec (Fig. 4); thus, they appear to be ancient Ecuadorian SP haplotypes and not SLC haplotypes.

All SLC populations, distributed from Mexico to southern Peru, were genetically close and composed mainly of hSL haplotypes. However, SLC MA was the most ancient SLC; its hSL haplotypes were the most diverse (Fig. 4) and exhibited the lowest LD (Fig. 5 C).

Passport data represent additional completely independent evidence in favor of the ancient origin of the SLC MA (Fig. S16). In Mexico, SLC has been collected mostly as a wild species in natural or disturbed environments and not under cultivation. Unfortunately, collectors were mainly interested in the degree of cultivation and did not differentiate between wild and ruderal environments. However, a recent study that involved an on-site evaluation reported wild SLC in tropical and mesophilic forests and shrublands (Álvarez-Hernández, Cortez-Madrigal, & García-Ruiz, 2009). This abundance of wild Mesoamerican SLC contrasted with its absence in the Andean region. In northern Peru and Ecuador, SLC was mainly cultivated or semi-cultivated in backyards. Semi-cultivated plants are those that, despite seldomly being planted, are cared for by backyard owners, as reported by Rick and Holle (1990). To our knowledge, Ecuadorian and northern Peruvian SLCs have never been reported in natural environments. Moreover, some Ecuadorian SLCs are commercially cultivated and have been mistakenly collected as vintage SLLs because they produce large fruits and are sold in markets (José Blanca et al., 2015). In southern Peru SLC, the ecology of SLC is different and is commonly found in disturbed environments. This SLC behavior has also been observed in most other subtropical areas worldwide. Thus, SLC has become a very successful invader in human-modified environments.

Mesoamerican and Andean hSL haplotypes have had no time to differentiate, indicating that the migration from Mesoamerica to Peru and Ecuador resulted from a rapid and recent event. Ruderal and weedy behavior could explain why SLC migrated back from Mesoamerica to Ecuador. SLC could have arrived in the Andean region from Mesoamerica as a weed associated with importing other crop species, such as Mesoamerican maize. Archeological records indicate that maize arrived in Ecuador approximately 7000–5500 calibrated years before present (Grobman et al., 2012; Meyer & Purugganan, 2013). If this hypothesis is true, it implies that the weedy SLC that arrived in the Andean region would have produced small fruits. The morphological characterization of the extant Mesoamerican SLC agrees with this hypothesis. Mesoamerican SLC typically produces small fruits. However, without direct archeological evidence, any hypotheses regarding the fruit phenotypes of populations thousands of years old are, by necessity, somewhat speculative, especially in populations that have undergone extensive migrations.

Moreover, haplotype composition, fastStructure, TreeMix, and ABBA-BABA results clearly showed that the Ecuadorian and Peruvian SLC populations were admixtures between SLC MA and SP, probably created after SLC MA migrated south to the Amazonian region and introgressed some Ecuadorian and Peruvian alleles. However, when the TreeMix analysis was conducted with less stringent LD thresholds (Fig. S10 B), it showed different tree topologies. Therefore, caution is advised when interpreting the TreeMix results. This lack of robustness is a limitation acknowledged by the TreeMix authors (Pickrell & Pritchard, 2012). Furthermore, it might have caused problems in the analysis conducted by Razifard et al. (2020) with an LD threshold that overrepresented the highly linked tomato pericentromeric regions.

Based on the high Nei diversity found in SLC Ec, all previous studies proposed SLC Ec and not SLC MA as the oldest SLC population (Jose Blanca et al., 2012; Lin et al., 2014; Razifard et al., 2020) (Fig. 9 B and C). However, this diversity index was calculated using all haplotypes, which was also found in the current study (Fig. 5 A and B). Moreover, the previous models struggled to explain how the highly diverse SLC Ec could have been derived from the genetically close but less diverse SP Ec, and they all proposed additional secondary contacts between SP and SLC to explain the high SLC Ec Nei diversity. However, this proposal undermined the main evidence used to appoint SLC Ec as the oldest SLC.

### There and back

The wide range of latitudes covered by the wild SP and SLC plants suggests that, in the northward migration of SP from Peru to Mexico, which became SLC, there should have been some selection related to latitudinal adaptation. Moreover, some of the adaptations associated with the northern latitudes could have been detrimental in Ecuador and might have been reverted in the southward trip of SLC to Ecuador. However, given that the SLC southbound migration was too fast for many new haplotypes to arise, we hypothesized that the alleles used in the readaptation of SLC to Ecuadorian latitudes would have been mainly introgressed from Ecuadorian SP. This hypothesis was successfully tested by calculating the ABBA-BABA statistics. The introgressions detected were concentrated in specific genomic regions (Fig 6 and Fig. S12). Moreover, many of these regions with SP introgressions have lower expected heterozygosity in SLC MA (Fig 6 C). This result indicated that many regions that suffered selective sweeps in the slow northward migrations recovered the original SP allele in the fast southward migration by introgression.

We manually inspected the genes located in these genomic regions (Table S3). Some regions might have been selected to adapt the plants to new lighting conditions. For example, in the region detected on chromosome 7, there are only five genes, and one is a FAR-like gene involved in light detection (Xie et al., 2020). In total, of the 13 analyzed regions, three included FAR-like genes, and one included a light response gene (Solyc02g021650). Other regions showed flowering-related genes: three had Agamous-like genes (Solyc06g161130, Solyc05g056620, Solyc04g160300), possibly involved in the regulation of flowering (Pan et al., 2010), and one had a possible flowering time regulation gene (Imamura et al., 2015). These regions also included a chloroplast development gene (Solyc06g050620) (Pérez-Pérez et al., 2013) and a photosystem protein (Solyc06g009950). Some of these regions are quite large, they include many genes, and it is impossible to know with certainty which gene was selected because most introgressed genes would have merely been carried over along with the selected ones. However, the abundance of the biological functions in these regions indicates that they are involved in latitudinal adaptations. The genetic studies required to evaluate these possibilities, gene by gene, might be accelerated by the public availability of hundreds of F2 populations of many of the accessions involved in the current study with SP, SLC, and SP parents (Mata-Nicolás et al., 2020).

This relationship between flowering genes and latitudinal adaptation has also been detected in other species. For instance, in potatoes, the southern wild species eased the introduction of cultivated potatoes to southern latitudes in Chile (Hardigan et al., 2017). Additionally, Cui et al. (Cui et al., 2020) observed the selection of genes related to heading date in the northward expansion of rice cultivation. In wheat and barley, photoperiod sensitivity arose when these crop species emerged from the Fertile Crescent (Meyer & Purugganan, 2013). Unfortunately, these resources are not available for all species. Concerning pumpkins and gourds, for instance, another American domesticate, they could not be transferred effectively between latitudes and were domesticated independently from different species in different regions (Kates, Soltis, & Soltis, 2017).

Therefore, the original wide range of latitudes covered by wild tomato plants might have created a wealth of allelic diversity that breeders could actively use to adapt tomato varieties to different latitudes worldwide. This might prove to be a key characteristic in the adaptation to climate change in the near future.

### Two-step domestication

Both Blanca et al. (2015) and Razifard et al. (2020) proposed a two-step domestication evolutionary model (Fig 9. B and C). According to these models, SLC would have been domesticated in northern Peru and then moved to Mesoamerica, where it was finally improved and transformed into SLL. The analyses and evidence shown in the current study are in agreement with this two-step domestication. The Mexican SLL is closely related to the Peruvian SLC. SLL Mx has few novel alleles and shares most of its haplotypes with SLC Pe, with some being SP Pe introgressions. Therefore, SLL Mx could result from improvements in local plants originally imported from northern Peru (Fig. 7). This close relationship between SLC Pe and SLL Mx constitutes indirect evidence in favor of Andean domestication. If there were semi-domesticated SLC in Mexico, Mexican growers would have probably derived SLL from it and not from imported Peruvian SLC.

Moreover, northern SLC Pe and SLC Ec, which, according to the proposed evolutionary model, are related to the oldest cultivated populations, were collected mainly from backyards, whereas in Mesoamerica, SLC was mainly wild and ruderal. Additionally, according to the morphological PCA (Fig. 8 A), the large-fruited SLC morphological type, typical of northern Peruvian SLC, was very close to the Mexican SLL, whereas the small-fruited SLC-type, typically found in SLC MA, was located between SP Ec and the large-fruited SLC. Thus, the sequence suggested by this morphological analysis would be as follows: SP Ec, small SLC (typical of Mesoamerican SLC), large-fruited SLC (typically found in northern Peru), and SLL. Therefore, there is a match between the evolutionary model suggested by morphological, passport, and genetic evidence. Most traits, such as style exertion, petal position and width, and fruit shape, varied monotonically along this sequence (Fig. 8 C). Assuming this progression, leaf type and margin would have already acquired its typical cultivated form in SLC MA; however, petal width would have decreased until its minimum was reached in SLC Pe N, and this population would also be the first one without folded back petals. Inflorescences gradually became more irregular starting with SLC MA and reached a maximum in SLL Mx, a trend shared by the flat and ribbed fruits that would have appeared in SLC MA and SLC Pe N, respectively. Razifard et al. (2020) noticed this same pattern of small-fruited SLC in Mesoamerica. However, because they thought Peruvian and Ecuadorian SLC to be older than Mesoamerican SLC, they proposed a reduction in fruit size during the migration out of the Andean region and a redomestication in Mesoamerica. The model presented in the current study proposes a smoother domestication process (Fig. 8), and it explains why the same domesticated alleles were found in Ecuador, Peru, and Mesoamerica (José Blanca et al., 2015).

### Conclusions

The new analysis method based on Procrustes and automatic haplotype classification allowed us to propose a new hypothesis for the complex evolution of wild and cultivated tomato plants. The wild populations were Peruvian and Ecuadorian SP, and Mesoamerican SLC. After migrating back to Ecuador and Peru, SLC was domesticated, and the Mexican SLL would have been derived from these improved materials. This model is backed by traditional population genetic indexes, parametric statistical models, morphological and passport data, and the new haplotypic analysis. We hope that similar approaches can be used to study the complex domestication histories of other species. Finally, we identified genomic regions associated with the latitudinal migration experienced by tomato plants that could be useful for adapting the currently cultivated varieties to new latitudes, particularly in a world affected by climate change.

## Supporting information

Supplemental Table 1. Passport and morphological data

Supplemental Table 2. Sample data and mapping statistics.

Supplemental Table 3. Introgressed and selected regions

## Acknowledgements

This research was supported by the National Natural Science Foundation of the USA Varitome Project (NSF IOS 1564366).

## Data accessibility

The new genome reads supporting the conclusions of this article are available in the SRA repository under bioproject PRJNA702633 (https://www.ncbi.nlm.nih.gov/bioproject/702633).

All the Python code used is available in the Github public repositories: tomato_haplotype_paper and variation5 (https://github.com/bioinfcomav/tomato_haplotype_paper, https://github.com/bioinfcomav/variation5) under the GNU GPL license.

## Author contributions

JB and JC wrote the manuscript and designed the methodology. DM, PZ, JMP, JB, and JC analyzed the data. MJD performed the morphological characterizations. JMP, MJD, and EK participated in the discussion and manuscript revisions. All the authors have read and approved the final manuscript.

