## Supplementary material for "Haplotype analyses reveal novel insights into tomato history and domestication including long-distance migrations and latitudinal adaptations"

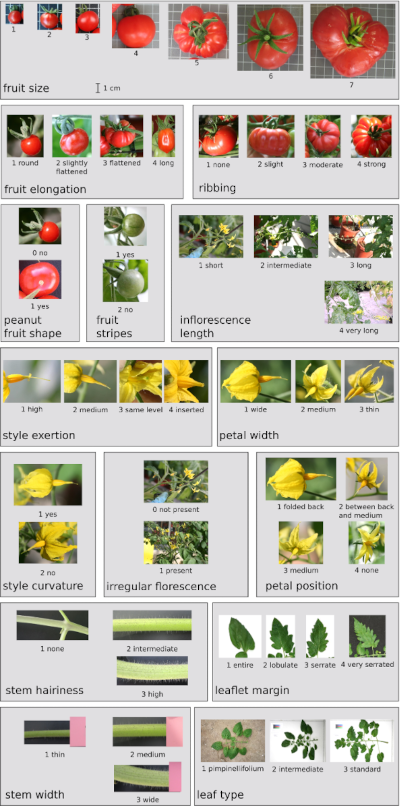

**Fig. S1.** Examples of the different morphological trait forms.

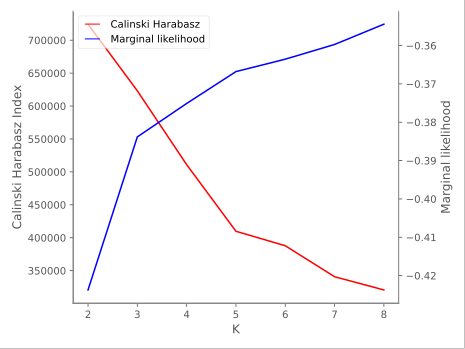

**Fig. S2**. FastSTRUCTURE marginal likelihoods and Calinski-Harabasz indexes calculated for different numbers of ancestral populations and haplotype types.

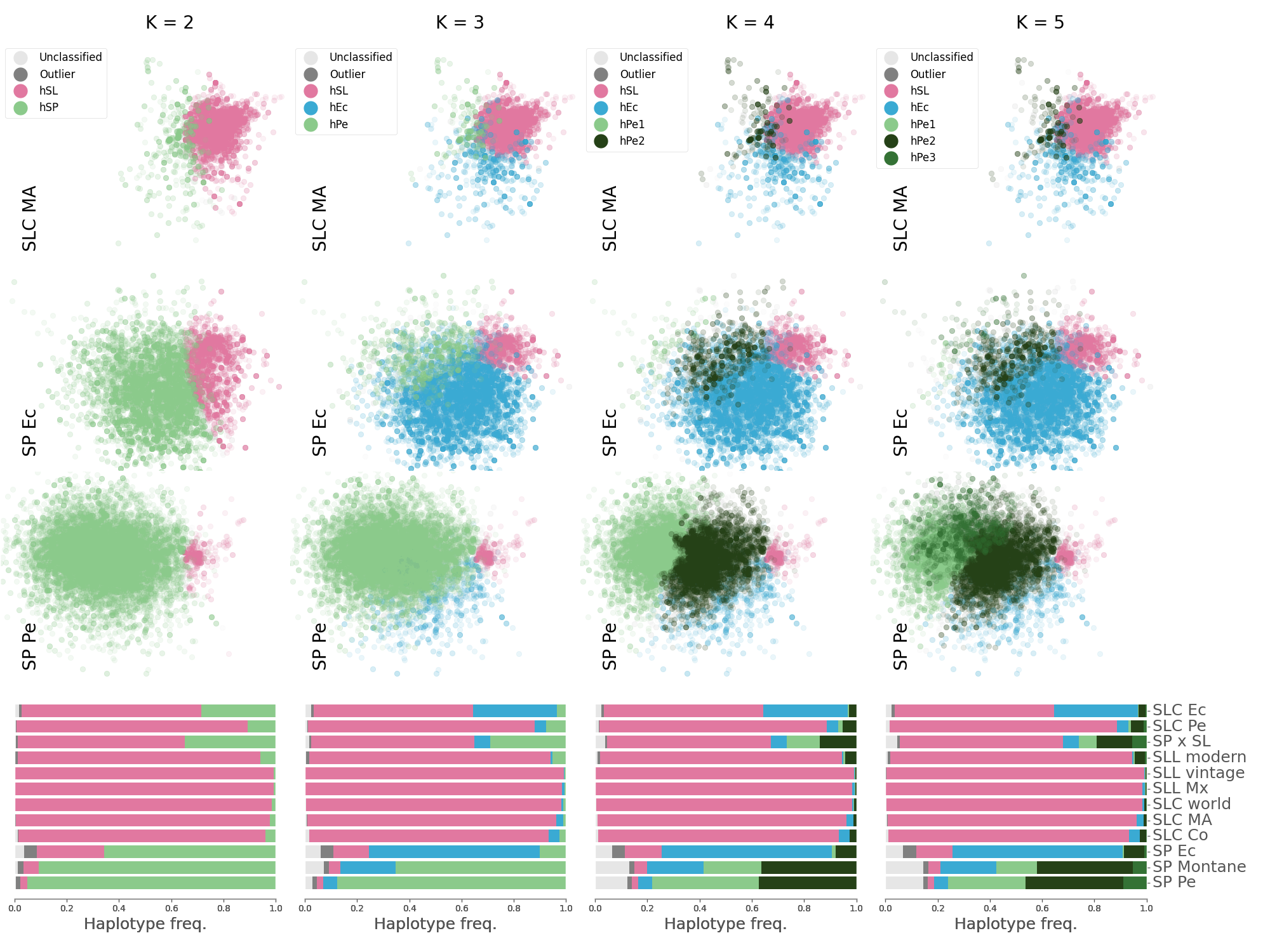

**Fig. S3.** Population haplotype composition calculated for different number of haplotype types. PCoA was carried out for every 500 kb genome segment using edit distances between haplotypes. The resulting PCoAs were aligned using Procrustes and automatically classified into three haplotype types. The classified haplotype PCoA was divided into several figures, one per population. In each figure only the haplotypes that belong to each population accessions were included. Frequencies of each haplotype type found in each population

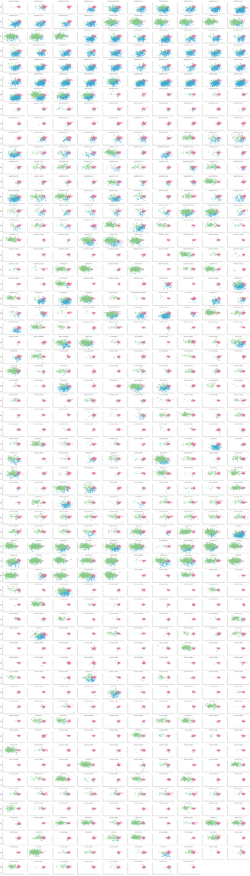

**Fig. S4.** Haplotype PCoA data of euchromatic regions aligned via Procrustes and automatically classified into three types plotted independently for every accession.

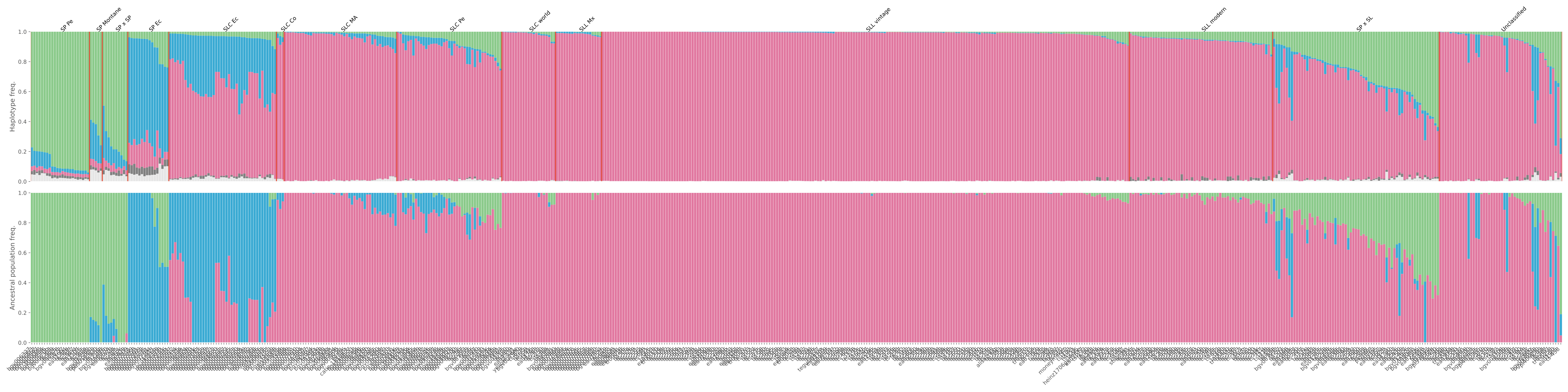

**Fig. S5.** Genome composition represented by the percentage of haplotype types, and by the fastSTRUCTURE ancestral population admixture proportions calculated for every accession.

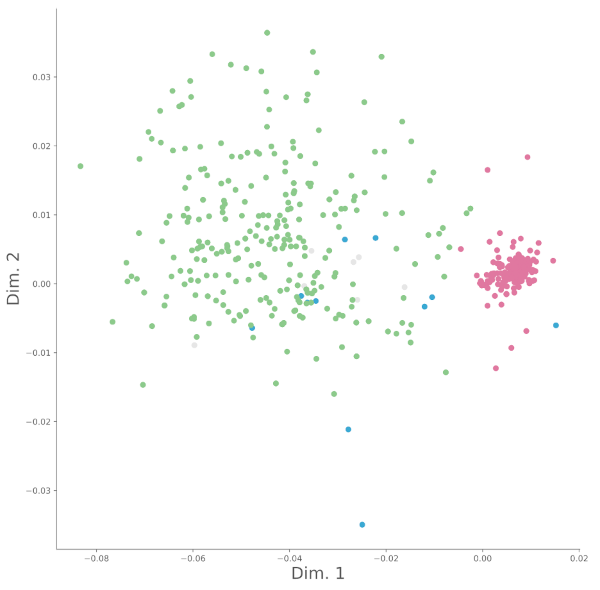

**Fig. S6.** Haplotype PCoA data of euchromatic regions aligned via Procrustes and automatically classified into three types plotted for the Cervil accession.

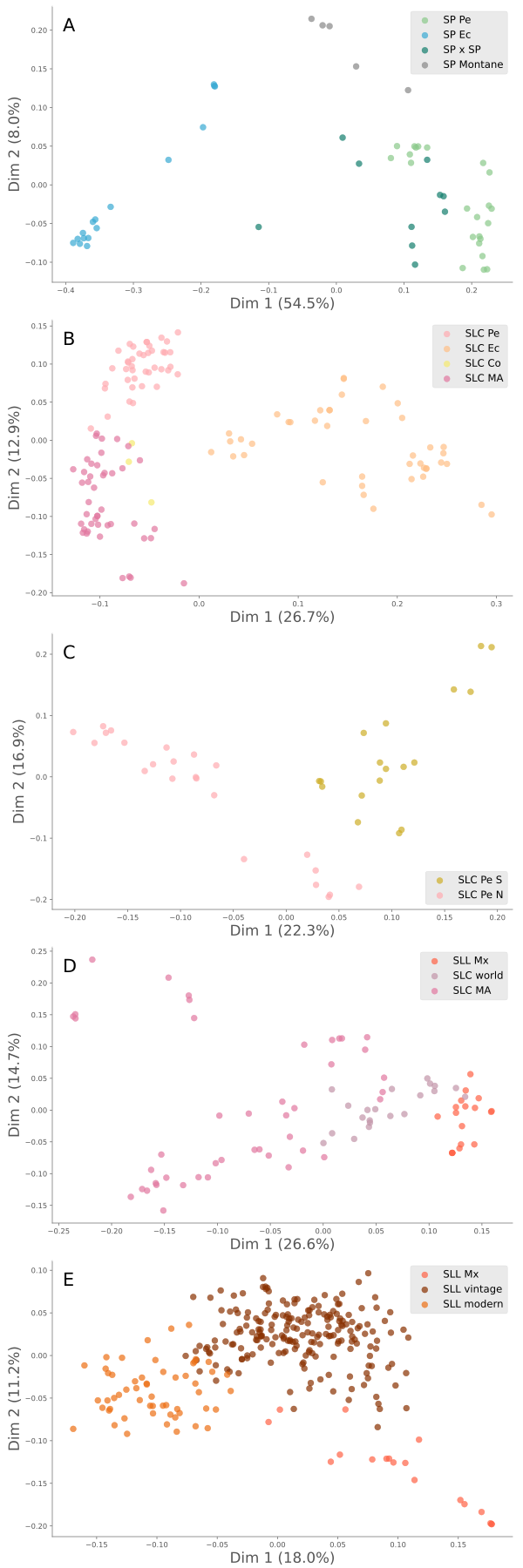

**Fig. S7.** Accessions based PCoA calculated from Kosman genetic distances for different populations calculated with variants characterized by a major allele frequency lower than 0.95 and a missing genotype rate lower than 0.1. Only the most variable variant from every 100 kb genomic segment was used. A) SP, B) SLC, C) Peruvian SLC, D) Mesoamerican SLC, and E) SLL.

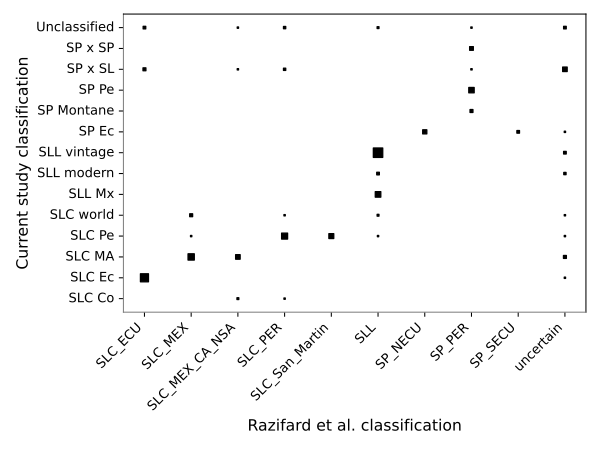

**Fig. S8.** Accession classification comparison between Razifard et al. (2020) and the current study. The marker size is proportional to the number of accessions.

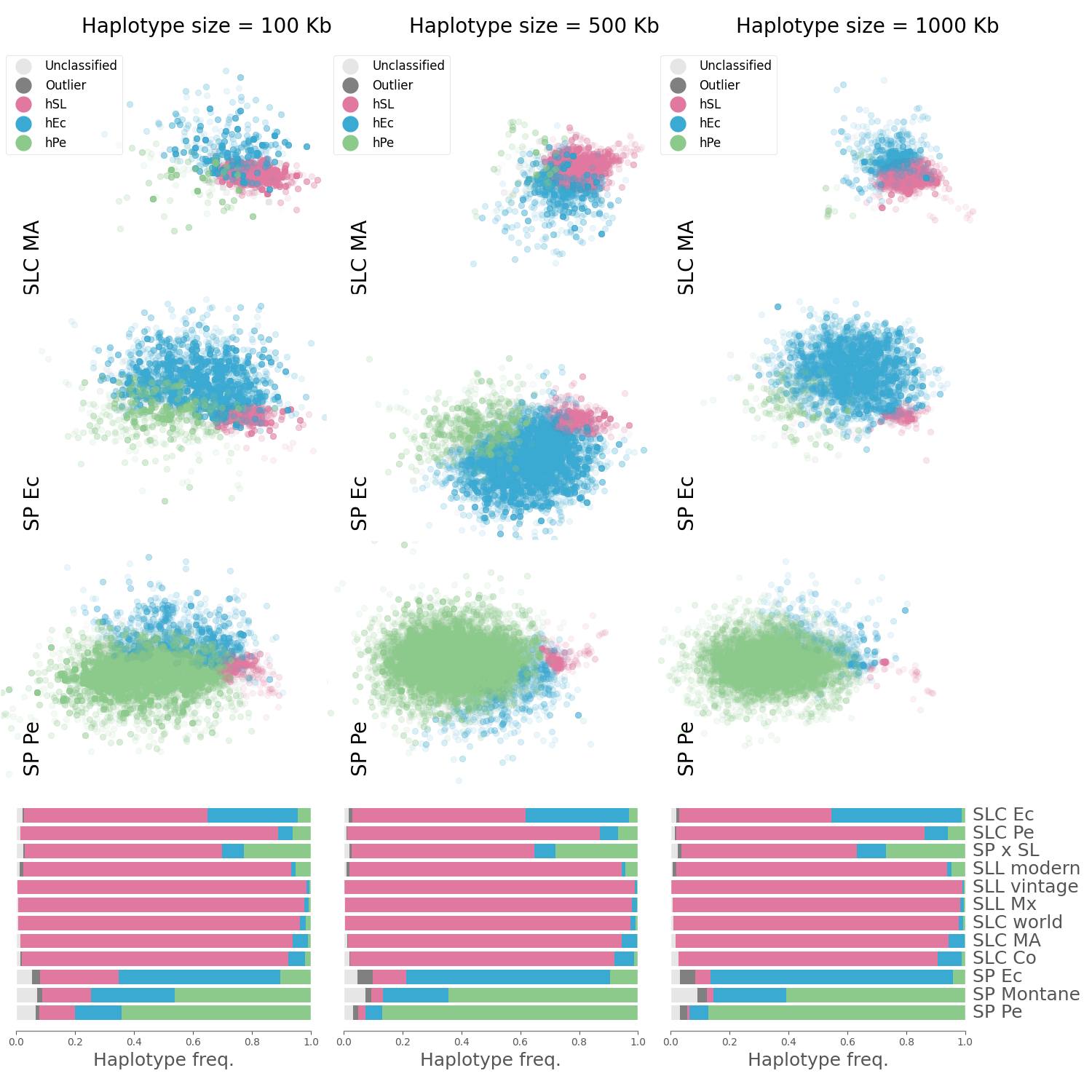

**Fig. S9.** Population haplotype composition calculated for different genome segment sizes: 100 kb, 500 kb, 1000 kb. PCoA was carried out using edit distances between haplotypes. The resulting PCoAs were aligned using Procrustes and automatically classified into three haplotype types. The classified haplotype PCoA was divided into several figures, one per population. In each figure only the haplotypes that belong to each population accessions were included. Frequencies of each haplotype type found in each population are shown in the bottom panel.

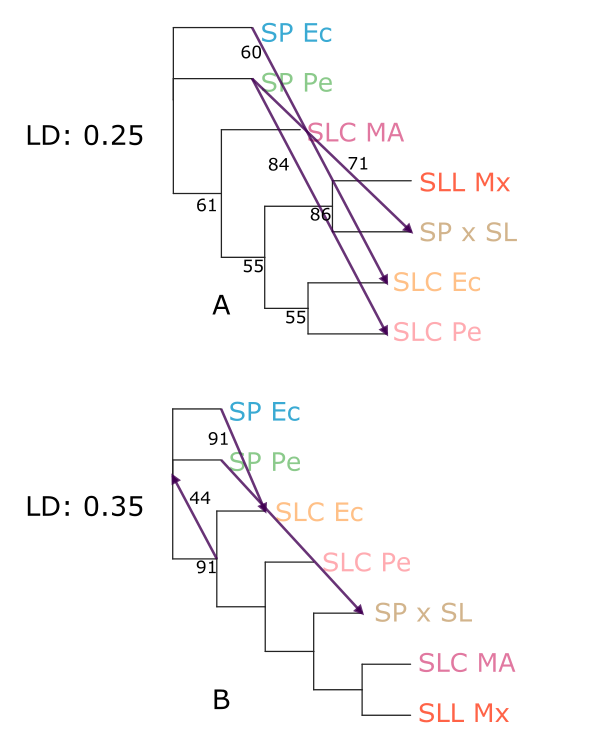

**Fig. S10.** TreeMix trees calculated for three migrations. The euchromatic regions were divided into blocks in which the variants had a linkage disequilibrium higher than the given threshold (0.25 for tree A and 0.35 for B). Only one variant per LD block was used to calculate the trees.

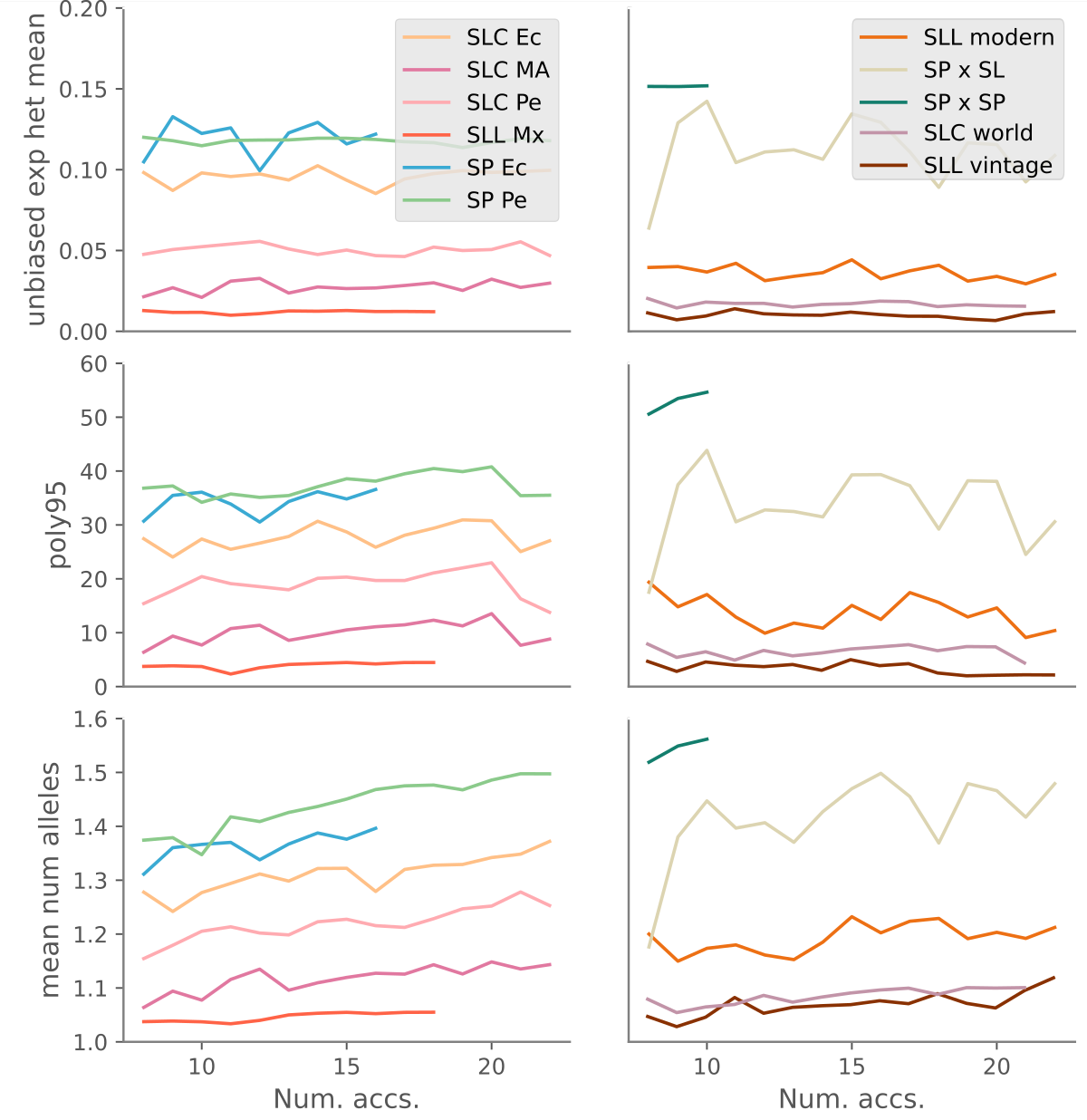

**Fig. S11.** Rarefaction analysis for three diversity indexes calculated for the genetic variants: unbiased expected heterozygosity, number of polymorphic variants (95% threshold), and mean number of alleles. The different populations were split into two panels to avoid excessive agglomeration of lines.

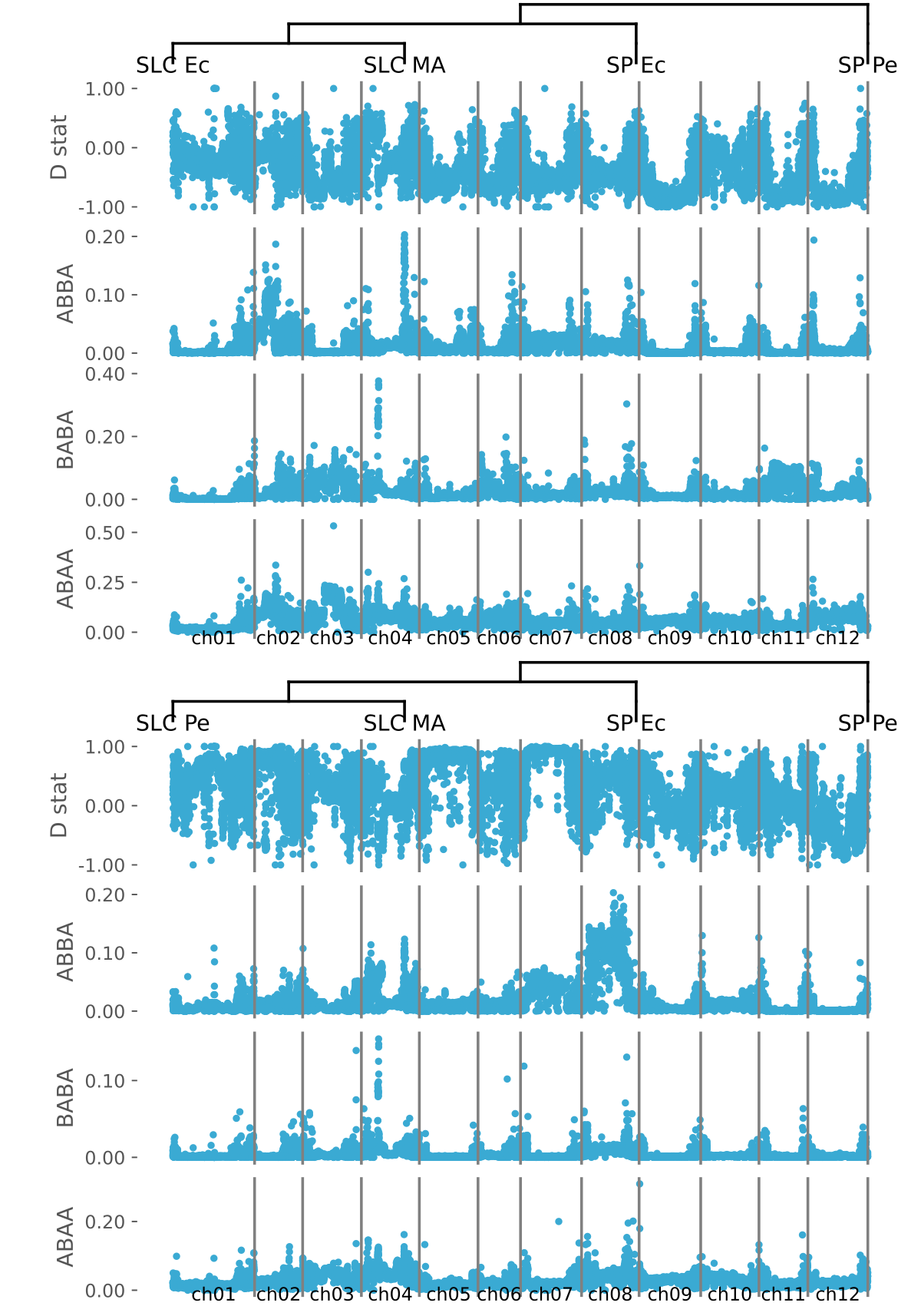

**Fig. S12.** D, ABBA, BABA, and ABAA products calculated in 500 kb segments along the genome using two different evolution models: SLC Ec, SLC MA, SP Ec, SP Pe and SLC Pe, SLC MA, SP Ec, SP Pe

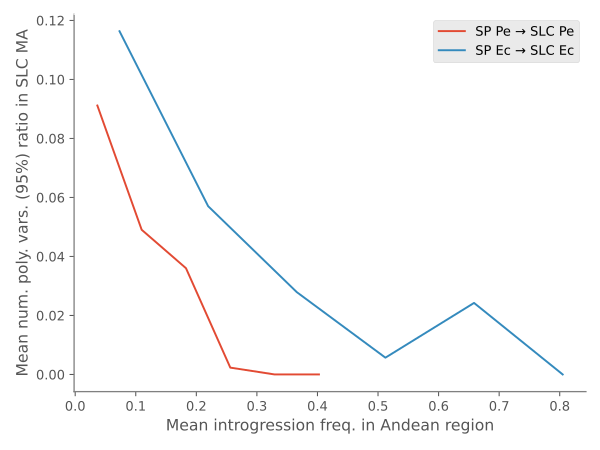

**Fig. S13.** Correlations between genomic diversity in the Mesoamerican SLC population and the introgressions from SP to SLC found in Peru and Ecuador. The Mesoamerican SLC diversity for each genomic segment (500 kb) was calculated as the number of polymorphic variants (95% threshold) found in the segment. An allele was considered to be introgressed from SP to SLC when it was not found in Mesoamerican SLC, which in this analysis was assumed to be the ancestral SLC population, but it had an abundance higher than 0.1 in SP.

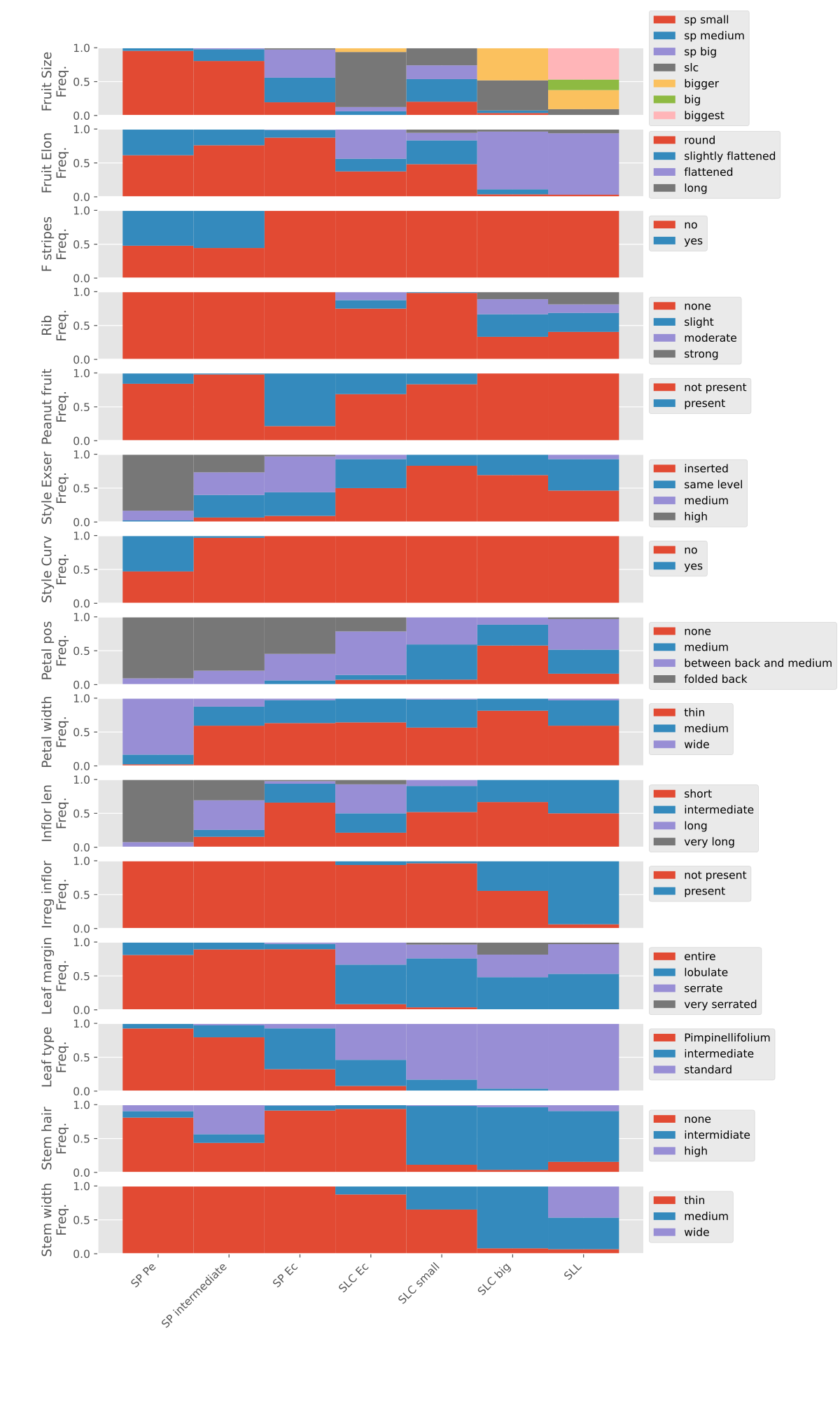

**Fig. S14.** Morphological group characterization. The frequency of each trait form in each morphological type is represented.

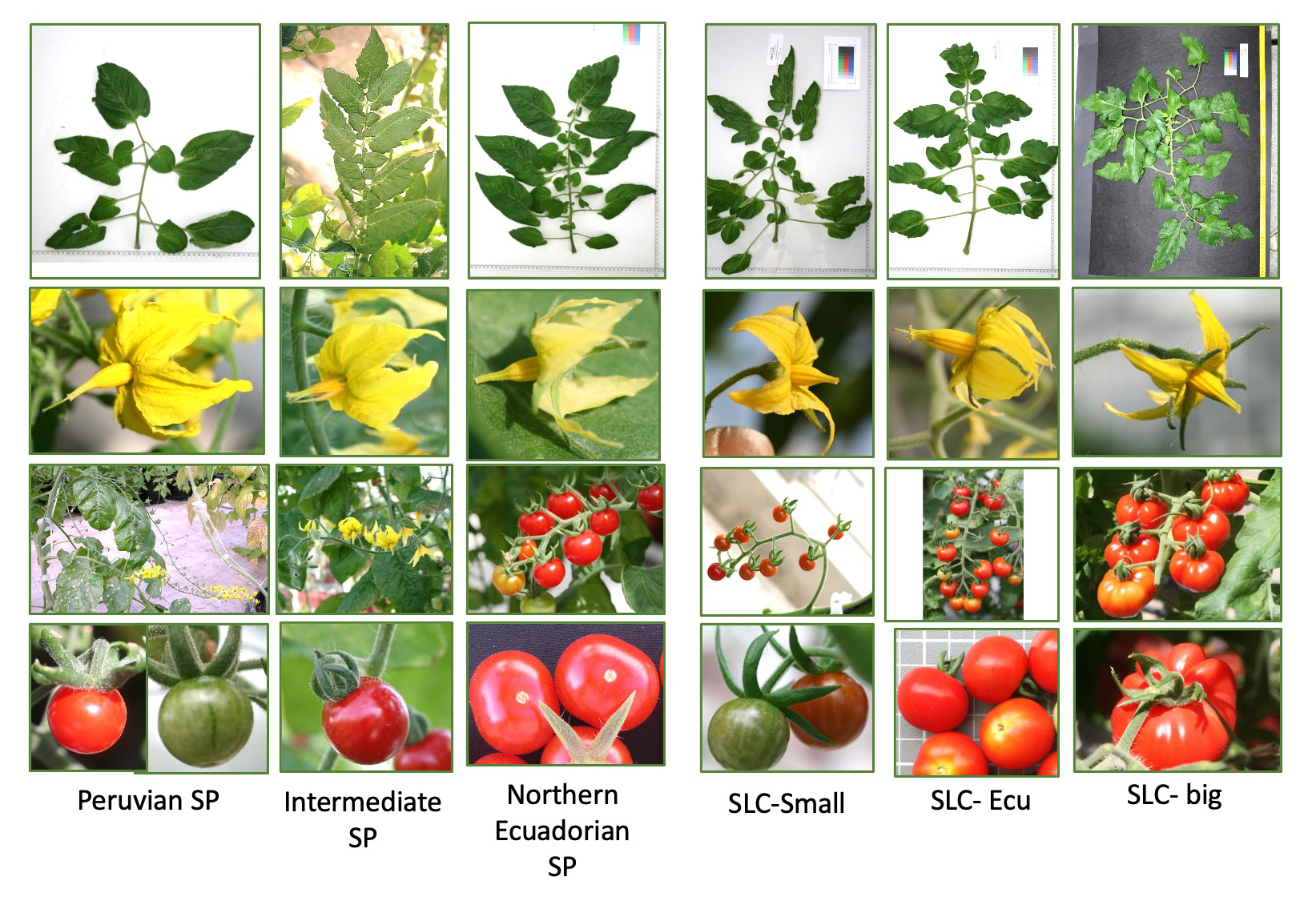

**Fig. S15.** Representative images of the different morphotypes.

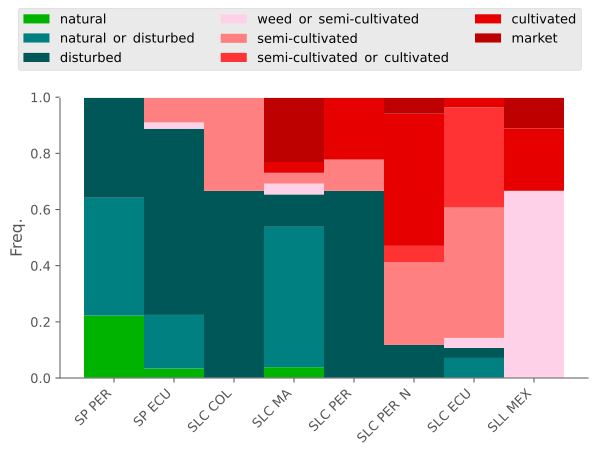

**Fig. S16.** Collection site characterization for the different populations taken from the passport data. Collection sites were classified in natural, disturbed, weed, semi-cultivated, and cultivated. The frequency of accessions collected in each type of environment for each genetic population is represented.

**Supplementary Table 1**. Accession origin, mean read coverage and genetic classification.

**Supplementary Table 2**. Annotation of genomic regions with a high frequency of introgression in SLC Ec, and SLC Pe and a low diversity in SLC MA.

**Supplementary Table 3**. Passport and morphological characterization data.
